# Sex-specific transcriptome similarity networks elucidate comorbidity relationships

**DOI:** 10.1101/2025.01.22.634077

**Authors:** Jon Sánchez-Valle, María Flores-Rodero, Felipe Xavier Costa, Jose Carbonell-Caballero, Iker Núñez-Carpintero, Rafael Tabarés-Seisdedos, Luis Mateus Rocha, Davide Cirillo, Alfonso Valencia

## Abstract

Humans present sex-driven biological differences. Consequently, the prevalence of analyzing specific diseases and comorbidities differs between the sexes, directly impacting patients’ management and treatment. Despite its relevance and the growing evidence of said differences across numerous diseases (with 4,370 PubMed results published within the past year), knowledge at the comorbidity level remains limited. In fact, to date, no study has attempted to identify the biological processes altered differently in women and men, promoting differences in comorbidities. To shed light on this problem, we analyze expression data for more than 100 diseases from public repositories, analyzing each sex independently. We calculate similarities between differential expression profiles by disease pairs and find that 13-16% of transcriptomically similar disease pairs are sex-specific. By comparing these results with epidemiological evidence, we recapitulate 53-60% of known comorbidities distinctly described for men and women, finding sex-specific transcriptomic similarities between sex-specific comorbid diseases. The analysis of shared underlying pathways shows that diseases can co-occur in men and women by altering alternative biological processes. Finally, we identify different drugs differentially associated with comorbid diseases depending on patients’ sex, highlighting the need to consider this relevant variable in the administration of drugs due to their possible influence on comorbidities.

## Introduction

Comorbidity and multimorbidity, defined as the presence of more than one medical condition in individuals ^1,2^, have been studied by conducting cohort, case-control, and nested case-control studies. Such studies have described, for example, that patients with type II diabetes (T2D) have a higher prevalence of dementia and cancer, with men specifically showing a higher risk of Alzheimer’s disease ^3,4^. However, the development of a digital medical record system ^5^ and the implementation of the Veterans Health Information System and Technology Architecture ^6^ paved the way for the collection of medical information in the so-called Electronic Health Records (EHR). Since then, several studies have systematically analyzed comorbidity relations and constructed networks based on epidemiological evidence. For example, Hidalgo *et al*. generated a network exploring Medicare data from more than 30 million patients in the United States ^7^. They found insightful differences in comorbidity relationships between genders, such as a higher risk of nephropathies in women or acute myocardial infarction in men with T2D. In addition, Jensen *et al*. generated chains of disease diagnoses - temporal disease trajectories - concatenating statistically significant co-occurrences using EHRs from Denmark, identifying key diseases that contribute to the progression of other diseases and increase mortality ^8^. In the same population, Westergaard *et al*. found that women present more comorbidities than men and a higher number of disease co-occurrences with an increased timespan between diagnoses ^9^. Therefore, those works suggest that men and women present essential differences in disease comorbidities, highlighting the importance of understanding such differences from the molecular point of view.

Since the publication of the first human disease network in 2007 by Goh *et al*., where diseases were connected if they shared at least one altered gene (diseasome) ^10^, dozens of works have studied molecular similarities between diseases, some of them trying to understand comorbidities better. Lee *et al*. described that diseases with coupled metabolic reactions co-occur 3 to 7 times more than those without coupling reactions ^11^. Measuring distances between disease modules in the interactome showed that diseases with overlapping modules tend to co-occur more than expected by chance ^12,13^. Transcriptomic similarities between diseases significantly recovered epidemiologically described co-occurrences ^14,15^. Recently, Dong and colleagues combined EHR and Genome-wide association studies (GWAS) data from the UK Biobank to recapitulate 46% of the multimorbidities ^16^.

We have already pointed out the lack of specific analysis of disease networks in terms of sex differences, despite its potential importance at the scientific and medical level ^17^, even more when we know that 37% of all genes exhibit sex-biased expression in at least one tissue ^18^. Liu *et al*. found disease-associated sex-specific polymorphisms in GWAS ^19^. Constructing sample-specific gene regulatory networks from healthy human tissues, Lopes-Ramos *et al*. found that many transcription factors have different regulatory targets depending on the sex of the sample. Interestingly, those differentially targeted genes are enriched in tissue-related functions and diseases. As an example, Alzheimer’s disease genes are targeted by different transcription factors depending on the sex of the sample ^20^.

Given the knowledge gap regarding sex differences at the comorbidity level, we have generated disease transcriptomic similarity networks separately for men and women. To our knowledge, this represents the first analysis using this approach in the field (see Figure 1). To avoid the constant change of terminology (female/male and women/men), throughout the text, we will always use women and men to refer both to transcriptomic data (female/male), and epidemiological data (women/men). The generated networks recover a representative set of known comorbidities previously described for women and men ^9^. Analyzing pathways altered in the same direction in comorbid diseases allows us to propose hypotheses for differences in disease co-occurrence between women and men. In addition, we find that disease pairs may co-occur more than expected by chance in both sexes through different biological processes. Finally, we translate the results to potentially related drugs, highlighting the relevance of understanding the scientific and medical implications of studying sex-specific differences in diseases and comorbidities.

**Figure 1.**
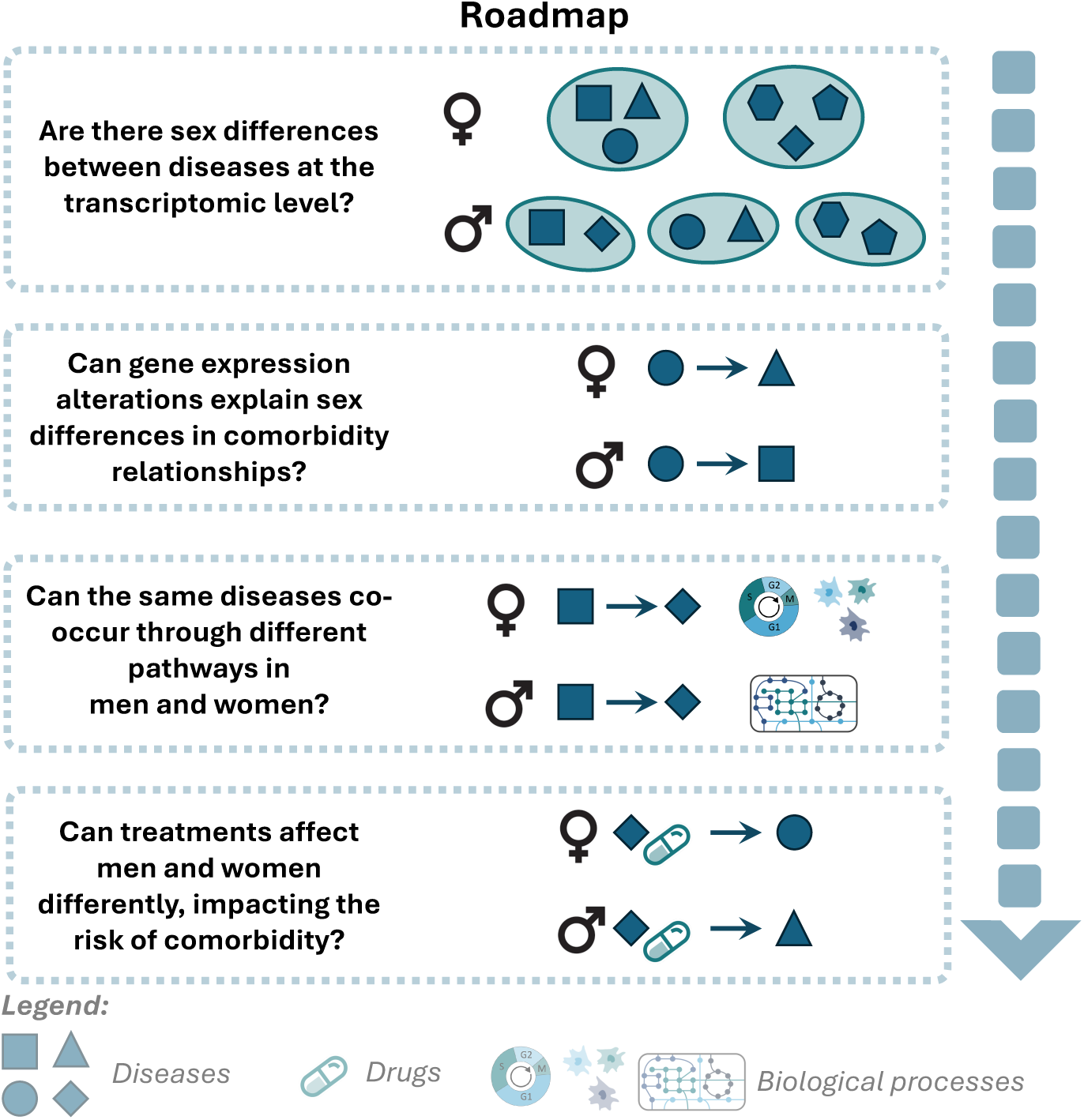
Study roadmap. Schematic representation of the main questions raised in the study. Squares, diamonds, triangles and circles denote diseases. Arrows indicate a higher-than-expected risk of developing a disease when suffering from a previous one.

## Methods

### Gene expression analysis

Raw expression data was downloaded from the Gene Expression Omnibus (GEO) and ArrayExpress (https://www.ebi.ac.uk/arrayexpress). Studies conducted on the *HG U133Plus 2 Affymetrix* microarray platform were chosen based on their cost-effectiveness and reproducibility, given their potential to translate findings into clinical practice. This selection also considered the large sample sizes necessary for analyzing disease-disease associations while minimizing biases that could arise from using different platforms or technologies. After removing low-quality samples (GNUSE median values higher than 1.25 ^21^), 128 diseases with at least three cases and three controls were kept. As samples’ sex information is only given for 52.23% (6,685/12,797) of the samples, the massiR package was used to infer samples’ sex, properly recovering the sex for 94.24% of the samples with annotated sex ^22^. This method classifies samples as male or female by analyzing the expression of the probes for Y chromosome genes. To homogenize the terms and to make the transcriptomic similarity networks that were generated in the following steps comparable to previously published epidemiological networks ^9^, the disease names were transformed into 3-digit codes of the International Classification of Diseases, version 10 (ICD10), of the World Health Organization, grouping specific diseases into a single 3-digit ICD code. As a result, we identified 76 (2,465 cases and 1,370 controls) and 77 (3,200 cases and 1,871 controls) ICD10 codes in women and men, 59 of which are common to both (see Supplementary Figure 1). 43% (42%) of the cases (controls) are women, with cases accounting for 63-64% of the samples in men and women, respectively. MAS 5.0 algorithm ^23^ was used to identify and remove lowly expressed genes (detection p-value<0.05). Background correction, normalization, and summarization were done using the frozen Robust Multiarray Analysis (fRMA) preprocessing algorithm (default parameters) ^24^. Differential expression analyses comparing the expression of samples with disease (cases) vs. disease-free samples (controls) were conducted using the linear regression model provided by the LIMMA package ^25^. These comparisons were performed separately and jointly for women and men, adjusting for confounding variables (study of origin and sex). Those genes with a False Discovery Rate (FDR) <= 0.05 and a log Fold Change (logFC) lower (higher) than zero were considered as significantly down- (up-) regulated.

### Gene set enrichment analysis

We performed functional enrichment analyses using the Gene Set Enrichment Analysis (GSEA) method ^26^, which was applied to the entire list of logFC-ranked genes obtained from the differential expression analysis (see previous section). The resources used in GSEA were Reactome, Gene Ontology, and KEGG. We performed disease clustering using Reactome pathways specifically, whose hierarchical structure allows us to identify 29 lowest-level pathway diagram annotations or categories. Pairwise Euclidean distances were calculated between Reactome pathway enrichment profiles (Normalized Enrichment Score values provided by GSEA). Ward2 method ^27^ was used to generate the clusters, identifying significant disease clusters through bootstrapping using the *pvclust* R package ^28^. The resulting dendrograms obtained for women and men were compared using *tanglegrams* from the *dendextend* R package ^29^, which gives two dendrograms (with the same set of labels), one facing the other, and having their labels connected by lines.

### Network construction

Transcriptional similarities were calculated using three distinct sets: (i) the complete list of annotated genes, (ii) the union of annotated genes with significant differential expression (sDEGs), and (iii) their intersection based on differential expression values (logFC) as previously done ^15^. Six similarity metrics were calculated: Pearson’s and Spearman’s coefficients, cosine similarity, and Euclidean, Canberra, and Manhattan distances. Empirical p-values were calculated through 10,000 permutations for the cosine similarity and the Euclidean, Canberra, and Manhattan distances, correcting for multiple testing by the Bonferroni approach and considering those similarities with an FDR<=0.05 as significant. In the case of Euclidean, Canberra, and Manhattan distances, the mean of the random distances was compared with the actual distances, obtaining positive (negative) values indicating a greater (lesser) similarity than expected by chance (see Supplementary Figure 2). The similarity values – obtained from the comparison between real and random distances in the case of Euclidean, Canberra and Manhattan distances, and from the coefficients in the case of Pearson and Spearman correlations and cosine similarity – were binarized, converting those coefficients greater than 0 to +1 and those less than 0 to -1. The results for the different metrics are similar, especially for Pearson and Spearman correlations, cosine similarity, and Euclidean distance (see Supplementary Figure 3). For simplicity, we will show the results obtained with Euclidean distances, which are the ones that allow us to recover more comorbidity relationships in a meaningful way (see Supplementary Table 1). Since the number of sDEGs strongly affects the ability to find similarities between diseases (correlations of 0.75 and 0.86 are obtained when working with the sDEGs union and intersection, compared to 0.34 when working with all genes, see Supplementary Figure 4), we focused on the calculations performed on the complete list of genes.

### Overlap with epidemiology

We used a previously published epidemiological network ^9^ (Supplementary Note 1) to identify the comorbidity relationships recovered by the disease transcriptomic similarity networks (DTSN) generated by comparing similarities between differential expression profiles and their ability to explain comorbidity relationships.

The overlap between networks was performed on the shared set of diseases (present in both the DTSNs and epidemiological networks). Specifically, the overlap of positive and negative transcriptomic similarities with the epidemiological networks was analyzed separately. Overlaps were measured by sex (women vs. women, men vs. men). The significance of the overlap was assessed by Fisher’s tests and the use of a set of specifically defined randomisations (generating 10,000 random networks shuffling the edges of the DTSNs while maintaining the degree distribution).

### Disease-drug associations

To study the potential sex-specific role of drugs in comorbidities, we retrieved drug targets from the DrugBank ^30^. Since the number of targets per drug is relatively small for enrichment analyses, we used the protein-protein interaction network extracted from IID ^31^ -selecting only those protein-protein interactions in humans that have been experimentally verified – to expand the number of targets associated with a given drug by mapping the targets on the network and selecting the first neighbours of the targets for each drug. We then conducted a GSEA ^26^ to associate drugs targeting the products of up or down-regulated genes with the corresponding disease, separately by sex. Disease-drug associations were extracted from the SIDER database ^32^. Disease names were transformed into ICD10 codes using the Unified Medical Language System ^33^, and DrugBank IDs were mapped into drug names.

## Results and discussion

### Sex-associated differences in gene expression

To examine the insights gained by studying the similarities between diseases at the transcriptional level according to the patient’s sex, we first downloaded microarray raw data from Gene Expression Omnibus (GEO, http://www.ncbi.nlm.nih.gov/geo) and ArrayExpress (https://www.ebi.ac.uk/arrayexpress). In order to study the insights gained by calculating similarities between diseases separately for women and men – rather than together – differential expression and enrichment analyses were performed comparing expression in cases vs. controls by analyzing both sexes together and individually (adjusting for sex differences; see methods).

To get a first overview of the differences in diseases between women and men, we performed hierarchical clustering on the Normalized Enrichment Score of their enriched pathways (see methods). On average, clusters were larger and more heterogeneous – comprising diseases from different categories – in women than men (66%/25% of the clusters in women/men were composed of diseases from different categories, see Figure 2). Indeed, the overall agreement between men and women clusters was quite limited. Parkinson’s and Alzheimer’s diseases (G20 and G30), previously linked at the molecular level ^34^, clustered significantly in men, sharing 112 pathways in which differentially expressed genes were enriched, mainly involved in signal transduction, immune system, and neuronal system. However, Alzheimer’s disease clustered with schizophrenia (F20) in women, sharing 116 pathways, including metabolic (glucose metabolism and respiratory electron transport) ^35,36^, metabolism of proteins (protein ubiquitination) ^37,38^ and neuronal system pathways (serotonin neurotransmitter release cycle and GABA synthesis release reuptake and degradation) ^39,40^, which have been previously associated with both diseases. Interestingly, a significant co-occurrence of dementia (including Alzheimer’s disease) has been described in patients with schizophrenia, with the risk being higher in women ^41^. In contrast, schizophrenia in men clustered with hepatitis C virus (B17) infection, two diseases that co-occur more than expected by chance in the population ^42^, with that risk being higher in men.

**Figure 2.**
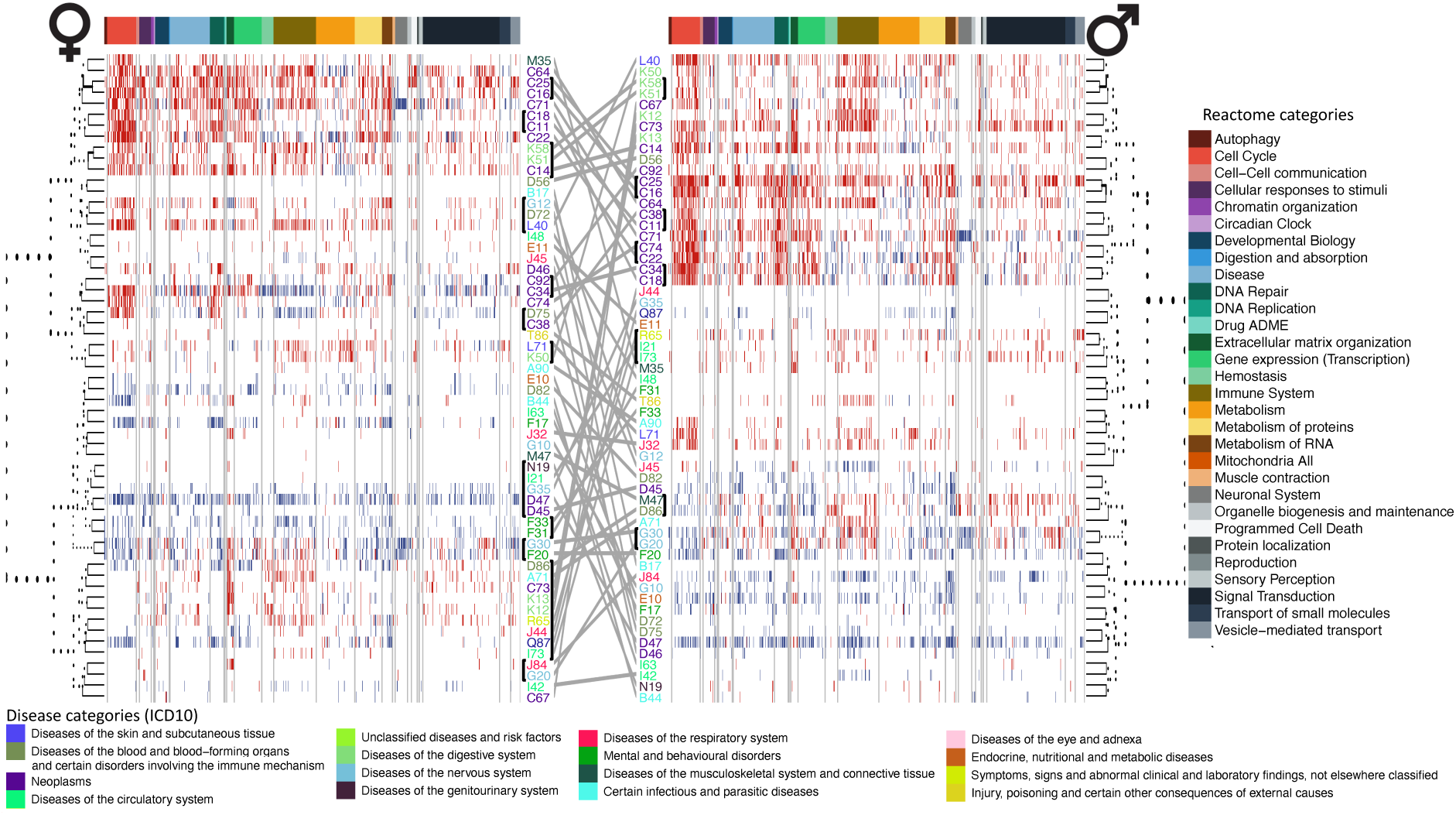
Reactome pathways significantly altered in disease in women and men. The diseases are represented in rows, and the pathways are in columns. The colors of the diseases refer to the category of the ICD10 to which they belong. The color bar in the columns represents the Reactome parents to which each pathway belongs. The red (blue) lines represent pathways that are under- (over-) expressed in each disease. The diseases have been clustered by calculating Euclidean distances between them and using the ward.D2 method on the Normalized Enrichment Scores. The gray lines connect diseases in women and men. Black lines near ICD codes indicate significant clusters after bootstrapping.

Only two pairs of diseases clustered together significantly in both sexes: pancreatic cancer - gastric cancer (C25 and C16) and irritable bowel syndrome (IBS) - ulcerative colitis (K58 and K51). Although both tumors clustered together in both sexes, cell cycle processes dominated the similarity in men and the immune system in women. Of particular note is the over-activation of interleukin signaling pathways, which has been proposed as a promising candidate for developing immunomodulatory therapeutic approaches ^43^. Indeed, considerable divergence in the tumor immune system has been described in tumors in women and men, highlighting its possible implications for differences in the efficacy of immunotherapy between sexes ^44^. In the case of the two digestive diseases (K58 and K51), they, in turn, clustered with oral cavity cancer (C14) in women, which co-occurs more than expected by chance, coinciding with the risk being higher in women ^45^. The increasing HPV prevalence amplifies the higher risk of oral cavity cancer in IBS patients and might be increased due to the use of immunosuppressive therapy ^46^, denoting the importance of treatments in comorbidity relationships. A total of 37 pathways were found to be altered in this cluster in women, 28 of them up-regulated (principally signal transduction pathways) and nine down-regulated (primarily metabolic pathways). Among them is the overactivation of NOTCH4 signaling, which has been proposed as a marker of oral cancer ^47^ ^a^nd whose expression is induced in macrophages after activation by Toll-like receptors (TLR) and interferon-γ (IFN-γ), which have been extensively studied in the context of IBS ^48^. These results highlight large differences at the molecular level between women and men in diseases belonging to different categories, opening the question of whether such differences may lead to the development of different comorbidities.

### Disease Transcriptomic Similarity Networks

Given the difference in how diseases cluster in women and men based on pathways, we built Disease Transcriptomic Similarity Networks (DTSN) by calculating similarities between pairs of diseases, adjusting for sex differences, and separately for women (wDTSN) and men (mDTSN) to study differences at the pairwise level. Similarities were calculated between diseases on the logFC (see methods).

Between 45% and 48% of all the potential edges in the wDTSN and mDTSN were positive, meaning the distance was smaller than expected at random. In contrast, 25-27% of the edges were negative, indicating larger-than-expected distances. As described in epidemiology ^9^, significantly more positive interactions were observed between diseases of the same category than between diseases of different categories in both women and men (Odds Ratio (OR)=2.81 in women vs. OR=1.83 in men, Supplementary Table 2), pointing out that transcriptional similarity may help to understand why diseases affecting the same system are more likely to co-occur versus those affecting different systems. These ORs were even larger when we considered only the connections that preserve the most similar comorbidity trajectories, i.e. the network backbone ^49^, (4.4 in women and 3.2 in men), denoting a greater relevance of intra-category connections in the flow of information, which is also observed in epidemiology (see Supplementary Note 2). Analyzing specific categories on mDTSN and wDTSN, diseases of the digestive and musculoskeletal system in men and skin diseases in women had a clustering coefficient of 1 (were all connected), followed by mental illnesses which, in women, had a density of 0.83 compared to 0.4 in men, correlated with the higher co-occurrence of mental diseases in women than in men ^50^ (see Supplementary Figure 5). Focusing on the set of diseases common to men and women, we found significantly fewer positive interactions in women than in men between the digestive system and nervous system diseases (OR= 0.21) and digestive system and blood diseases (OR= 0.27, see Supplementary Table 3). These results correlate with the higher risk of co-occurrence of IBS and multiple sclerosis ^51^, Parkinson’s disease ^52^, or dementia ^53^. In fact, no positive interactions were found between IBS and multiple sclerosis in men. In contrast, women had significantly more negative interactions between digestive and nervous system diseases (OR = 10.81, see Supplementary Table 4). When comparing the networks generated considering both sexes together and separately, 16.22% and 13.49% of the positive interactions were detected only in women and men, respectively, values that hover around 9% and 4% in the epidemiology ^9^ ^(^see Supplementary Figure 6A-B). These results demonstrate that, as in epidemiology, relationships relevant to only one of the sexes can be lost by analysing women and men together.

### Biological clues to differences in disease co-occurrences in men and women

Previous studies have shown that transcriptional similarity can significantly recover comorbidity relationships ^14,15^, even when analyzing very different populations, highlighting the relevance of the approach (see Supplementary Note 3). As reflected in the introduction, to date, this phenomenon has not been evaluated separately in women and men ^17^. Our results showed that 53.12% and 60.68% of the disease co-occurrences described in women and men by Westergaard et al. ^9^ ^w^ere also connected in the wDTSN and mDTSN (see Figure 3, Supplementary Figure 6C and Supplementary Table 1). Metastases and myelodysplastic syndromes were the ICD10 codes with the highest number of comorbidities retrieved by the wDTSN and mDTSN respectively (92.3% and 87.5% respectively, see Figure 3 and Supplementary Figure 7), where the former co-occurred only with other neoplasms, while the latter also co-occurred with blood disorders (autosomal dominant monocytopenia and primary myelofibrosis). As expected, pathways such as positive regulation of chromosome segregation, DNA unwinding involved in DNA replication, deposition of new CENP-A containing nucleosomes at the centromere, and DNA strand elongation were found to be up-regulated in eleven of the twelve cancers co-occurring with metastases (including women-specific cancers such as ovary and breast cancer). The role of these pathways in cancer has been extensively described ^54^, and they have been shown to correlate with tumor stage and be enriched in relapsed and metastatic tumor patients ^55^ ^a^nd to promote epithelial-mesenchymal transition (precursor of metastasis) ^56^. In the case of myelodysplastic syndromes, the same biological processes were found to be altered in the same direction in up to five of the seven recovered comorbidities. This is the case of down-regulation of T cell receptor signaling, altered in juvenile myelomonocytic leukemia (C93), primary myelofibrosis (D75), autosomal dominant monocytopenia (D72), neoplasms of uncertain behavior of lymphoid, hematopoietic and related tissue (D47) and polycythemia vera (D45). Immune deregulation has been shown to play a prominent role in the pathogenesis and progression of the mentioned disorders ^57,58^, opening the door to a possible role also in their co-occurrence. Indeed, there is a syndrome called MonoMAC, characterized by severe monocytopenia and a high risk of progression to myelodysplasia ^59^. Other additional processes related to the immune system, such as antigen receptor-mediated signaling, B-cell receptor signaling, and primary immunodeficiency, were also jointly altered in the last four. Genes such as JAK1 werer also found to be changed in expression in most diseases that co-occurred with myelodysplastic syndromes (four out of seven). The role of JAKs has been widely described in immunity, immunodeficiency, and cancer ^60^. Likewise, it is known that interleukin-7 is crucial for lymphocyte development and for the survival, proliferation, differentiation, and activity of mature T- and B-cells ^61^. Four of the seven diseases comorbid with myelodysplastic syndromes had altered IL7R levels.

**Figure 3.**
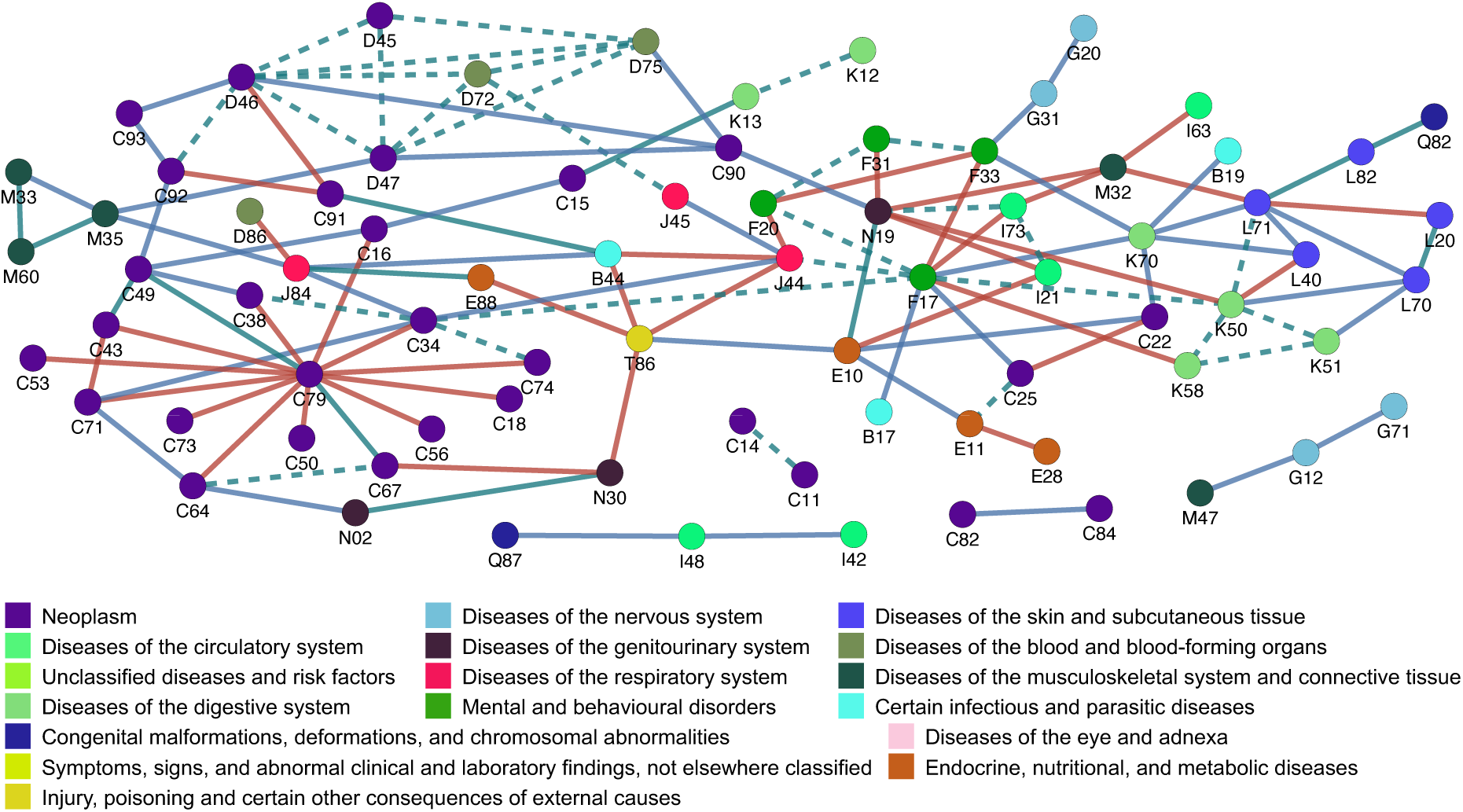
Comorbidities explained by the Disease Transcriptomic Similarity Network. Network of comorbidities recovered based on transcriptomic similarities. Each node represents a disease, colored based on the disease category they belong to (ICD10). Red, blue, and green edges represent comorbidities (retrieved by analyzing the epidemiology separately for women and men and adjusting for sex) recovered by calculating similarities at the gene expression level between diseases in the same way (separately for women and men and adjusting for sex differences). The dashed edges represent comorbidities described in women and men that are recovered by analyzing transcriptomic similarities separately for women and men.

In addition to co-occurrences of diseases of the same category, sex-specific DTSNs also identified altered biological processes in comorbid diseases of different categories. For example, interstitial cystitis (N30) has been shown to increase the risk of bladder cancer (C67) ^62^, being the risk only significant for women in the Danish population (RR=1.138, 97.5% CI: 1.032, 1.252). Nonsense-mediated decay, seleno-amino acid metabolism, and EIF2AK4 (GCN2) to amino acid deficiency pathways were down-regulated in both diseases in the wDTSN. Several studies have shown that inflammation plays a prominent role in interstitial cystitis ^63^. Interestingly, it has been described how upregulation of nonsense-mediated decay during chronic inflammation can help restore immune homeostasis ^64^, and in turn mutations that can elicit nonsense-mediated decay have been detected in half of patients with bladder cancer ^65^. In the same direction, oxidative stress is implicated in the pathogenesis of interstitial cystitis ^66^, and seleno-amino acids and their metabolic reprogramming regulate antioxidant defense, enzyme activity, and tumorigenesis ^67^, highlighting the widely known phenomenon that chronic inflammation and the resulting oxidative stress can increase the risk of cancer development ^68^.

Similarly, psoriasis was found to increase the risk of alcoholic hepatitis ^69^, two diseases positively connected in our mDTSN. Psoriasis is a clinically heterogeneous lifelong skin disease mediated by chronic inflammation ^70^, and alcoholic hepatitis is characterized by the degeneration of hepatocytes, neutrophil infiltration, and fibrosis ^71^. Previous studies have described that their co-occurrence might be driven by shared risk factors such as smoking and alcohol consumption, shared pathophysiological pathways including insulin resistance and oxidative stress, as well as psoriasis drugs potentially inducing liver damage ^72,73^. The mDTSN showed that most altered biological processes in both diseases implicated the cell cycle, signal transduction, and the immune system. Notably, several of these pathways were involved in the activation and signaling of nuclear regulatory factor κB (NFκB), which is a crucial mediator of the pathogenesis of psoriasis ^74^ ^a^nd at the same time is activated by chronic alcohol intake, promoting the hepatic macrophage expression of pro-inflammatory mediators ^75^ ^(^including the tumor necrosis factor involved in hepatocyte injury ^76^ ^a^nd psoriasis ^77)^. Supporting the existence of common pathways that could explain the co-occurrence of both diseases, it has been described that anti-TNF drugs can induce the onset of psoriasis ^78^ ^a^nd, in turn, significantly increase mortality in patients with moderate to severe alcoholic hepatitis ^79^. These diseases also co-occur more than expected in the other sex. Unfortunately, the lack of sufficient samples (see methods) has not allowed us to analyze these relationships.

Focusing now on the common set of diseases for which we have sufficient samples in both women and men, it has been described that patients with schizophrenia (F20) are at a higher risk of developing COPD (J44) ^80^, even when compared to smoking controls ^81^. At the core of such co-occurrence of diseases, it has been shown that patients with schizophrenia have impaired lung function ^82^. Interestingly, lung function SNPs acting as eQTLs have been linked to neuropsychiatric diseases, including schizophrenia ^83^. Clinical observations have shown that there are sex differences in prevalence, symptoms, and response to treatment in patients with schizophrenia ^84^, as well as in comorbidity relationships, where, for example, the risk of developing COPD would be higher in women ^69^ – indeed, in the Danish population the risk is only significant for women ^9^. The wDTSN recovered this comorbidity relationship between both diseases in women, finding pathways associated with mitochondria (mitochondrial translation, mitochondrial RNA metabolic process, and mitochondrial gene expression) and the immune system (interleukin 10 signaling, neutrophil degranulation, macrophage cytokine production, and antimicrobial peptides) altered in the same direction. Interestingly, previous studies have described an enrichment in mitochondrial processes in women vs men in schizophrenia ^85^ ^a^nd COPD ^86^, as well as differences in the immune system ^87,88^, supporting the hypothesis that differences between women and men at the disease level may explain the development of different comorbidities.

Similarly, a higher risk of developing IBS (K58) in smokers (F17) has been observed, being higher in women ^89^. Interestingly, the impact of smoking is greater in women than in men ^90^ ^a^nd the prevalence of IBS is higher in women ^91^, potentially driven by hormonal differences between both sexes ^92^. As in other comorbidities, most pathways in common detected in the wDTSN involved mitochondria (mitochondrial respiratory chain complex assembly, respiratory electron transport, complex I biogenesis, detoxification of reactive oxygen species) and immune system (neutrophil activation involved in immune response and immunoregulatory interactions between a lymphoid and non-lymphoid cell). Previous studies support the identified differences in each of the diseases separately ^93–95^, pointing to these processes as possible future targets for reducing the risk of disease co-occurrence. Other diseases that co-occur more than expected by chance in women but not men and were uniquely connected in the wDTSN include type I diabetes (T1D, E10) - myocardial infarction (I21) ^96^, bipolar disorder (F31) – uremia (N19), and COPD (J44) - chronic lung allograft dysfunction (B44, see Figure 3).

On the other hand, a higher-than-expected risk of developing liver cancer (C22) in men with T1D (E10) has been described ^97^. Interestingly, the prevalence of T1D is larger in prepubertal women and afterwards occurs twice as much in men ^98^. Most of the biological processes altered in the same direction in men in both diseases included mainly pathways associated with metabolism (metabolism of amino acids and derivatives, metabolism of vitamins and cofactors, glutathione metabolic process, reactive oxygen species metabolic process, and response to starvation) and the immune system (humoral immune response, positive regulation of immune effector process, and regulation of t helper 1 type immune response). Finally, by calculating similarities between diseases separately for each Reactome parent category (see Supplementary Note 4) we observed that gene expression and the immune system were the categories that recovered the most comorbidities (41-47%), and drug ADME the least (4-5%, see Supplementary Table 5). Despite the diversity in the amount of comorbidities identified for the different categories, all overlaps with the epidemiology were significant (see Supplementary Figure 8A-B and Supplementary Table 5). All the results presented in this section provide evidence of differences in the biological processes altered in diseases according to the sex of the patients and that these differences may help to understand better why disease co-occurrences are different between women and men. Among the most relevant biological processes, we find the immune system, metabolism, and mitochondria-associated processes as those that best explain the differences in comorbidity relationships between women and men, previously described to drive differences between _sexes 99,100._

### Comorbidities occur through different mechanisms in women and men

Once verified that differences in transcriptional similarities between diseases in women and men can explain differences in comorbidity relationships, the question is, is it possible that the mechanisms by which some diseases co-occur are different for each sex? This has been previously described for mechanical pain hypersensitivity, which is mediated through different immune cells in male (microglia) and female (T-cell) mice ^101^. As shown in Figure 4, 29 disease pairs co-occur in women and men more than expected by chance (12 belong to different categories), and such comorbidities are recovered in both wDTSN and mDTSN. However, there are sex-specific biological alterations.

**Figure 4.**
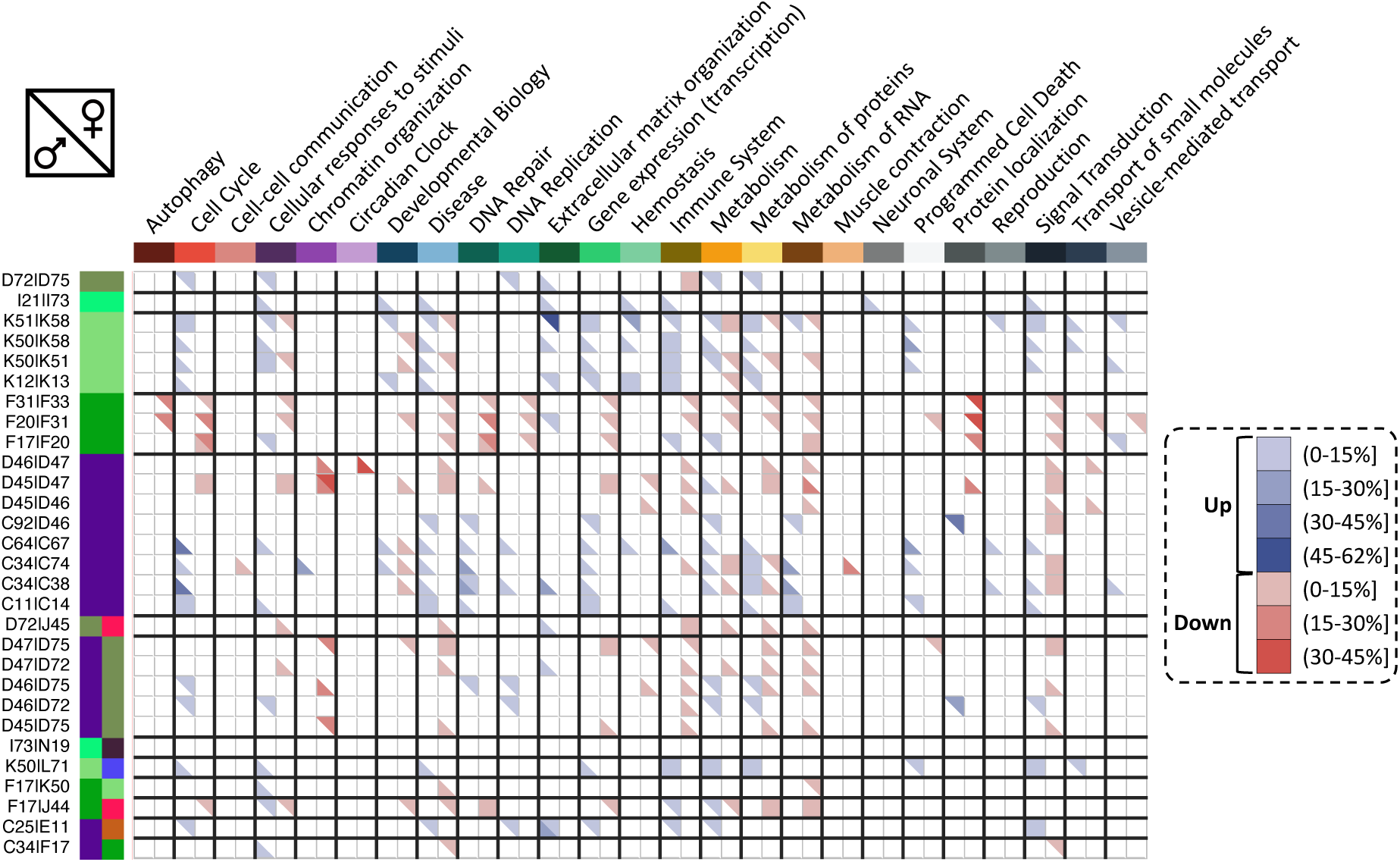
Pathways shared between comorbid diseases in women or men only. Heatmap with disease pairs in rows and Reactome categories in columns representing the percentages of pathways in each category that are up- (blue) or down-regulated (red) in both diseases. Each square is divided into two triangles, the one on the top (bottom) refers to women (men).

Smoking (F17) is a widely described risk factor for the development of COPD (J44), affecting both women and men ^102^. However, the mechanisms by which smoking increases the risk of COPD may differ by sex. As shown in Figure 4, in women, alterations common to both diseases were identified in the cell cycle, response to stimuli, metabolism, immune system, and developmental biology that did not occur in men; in men, alterations were observed in DNA repair and protein and RNA metabolism that did not occur in women. It has been previously proposed that cigarette metabolism may vary due to sex differences in the expression and activity of cytochrome P450 enzymes ^103^, a process that was found to be overexpressed in women in smokers and COPD but not in men (cytochrome p450 arranged by substrate type). As previously described, mitochondrial functional pathways were altered in both diseases in women but not men, which might be potential drivers of sex differences in lung diseases ^86^. In contrast, the processes of sumoylation of transcription cofactors, processing of capper intro containing pre-mRNA, and base excision repair were downregulated in men in both diseases but not in women.

Another example is the significant co-occurrence of pancreatic cancer (C25) and T2D (E11)^104^. It is interesting to note that men and women with pancreatic cancer share a high number of altered pathways in the same direction (Jaccard indexes for up and down-regulated pathways of 0.78 and 0.58, respectively), while the overlaps were low in the case of T2D (0.03 and 0, see Supplementary Table 6). These results point to greater sex differences in T2D than in pancreatic cancer. Therefore, the development of pancreatic cancer may be different according to sex in patients with T2D due to differences in its development ^105^. In men, T2D and pancreatic cancer shared mainly extracellular matrix organization pathways (collagen biosynthesis and modifying enzymes, laminin interactions, ECM proteoglycans…) and signal transduction pathways altered in the same direction (not shared in women). On the other hand, women shared immune system (neutrophil degranulation, antiviral mechanism by IFN stimulated genes…) and metabolism pathways (sphingolipid de novo biosynthesis and regulation of cholesterol biosynthesis by SREBP (SREBF)). The results highlight the usefulness of the proposed approach in identifying biological processes that are differentially altered in women and men and may be fundamental in the development of diseases and comorbidities differently in both sexes. Unfortunately, there is still a considerable gap in biological knowledge regarding the differences between women and men in the development of diseases, which is fundamental to understanding the differences in their comorbidities better.

### Sex differences in drug effects

In previous studies, we have identified drugs that could play a role in disease co-occurrence, both increasing and decreasing the risk, proposing this approach as a possible avenue for drug repositioning ^106^. Going a step further, we have identified subgroups of patients with a disease for which a specific drug association might be related to the observed increased risk of developing secondary diseases ^14^. In this instance, we investigate how different drugs may be associated with diseases differently in women and men and thus explain their specific comorbidity relationships. Due to the lack of studies analyzing differences in expression changes after drug intake as a function of sex, we extracted drug targets from DrugBank ^30^, expanded the association using a protein-protein interaction network ^31^, and performed enrichment analyses ^26^ to identify drugs whose expanded targets tend to be over- or under-expressed in different diseases as a function of sex. 3,878 DrugBank IDs were significantly enriched in at least one disease. Of them, 616 had associations (3,997) to 568 ICD10 codes in the SIDER database (a database of drugs and adverse drug reactions which contains drug indications, extracted from the package inserts using Natural Language Processing) ^32^. Focusing on the diseases for which there were sufficient samples in women and men, 43 of the 59 had at least one enriched drug associated with any diseases under analysis in SIDER. Studying the top 10 drugs associated with more diseases in women and men, only three were found in common (metformin, clofarabine and irinotecan), drugs known to treat diabetes, acute lymphoblastic leukemia, and metastasis. Arginine, bortezomib, and carfilzomib in women, and dexrazoxane, etoposide, and idarubicin in men were other drugs associated with more than 25% of the diseases (used to treat high blood pressure, cardiomyopathies, and cancer). Seven diseases did not share a single drug associated with both men and women: Aspergillus colonization of lung allograft, T1D, amyotrophic lateral sclerosis, interstitial lung disease, rosacea, connective tissue disorders, and axial spondyloarthropathy (see Supplementary Figure 9), while 8 presented a Jaccard index above 0.5: pancreatic, colon, and kidney cancer, neoplasms of uncertain behavior of lymphoid, hematopoietic and related tissue, oral dysplasia, Job’s syndrome, thalassemia and IBS, highlighting that there was great heterogeneity in the number of drugs shared between sexes depending on the disease. In total, women had more drugs associated with them in 19 diseases versus 20 diseases in which more drugs were associated with men. These differences between sexes were not due to differences in the number of samples of each sex (p-value of the correlation = 0.64). In fact, in diseases such as schizophrenia, 3.13 more drugs were identified in women than in men, while there were 2.7 more men than women. When analyzing the drugs enriched for diseases in women and men (and the diseases for which they are indicated), we found interesting differences according to sex (see Figure 5 and Supplementary Figure 10).

**Figure 5.**
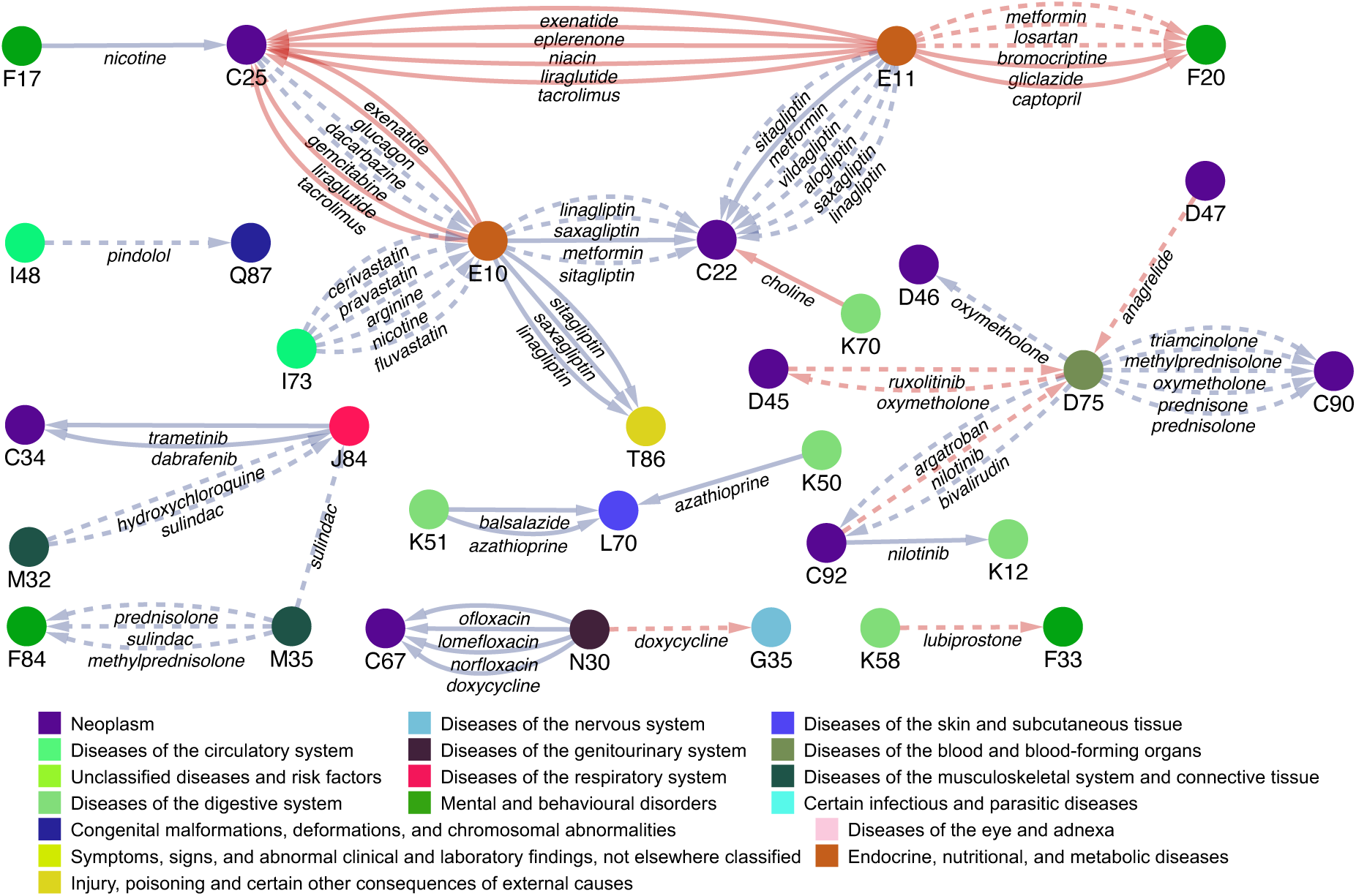
Sex-specific disease-drug associations. The nodes represent diseases, colored according to the disease category (ICD10) to which they belong. An edge connects two diseases if the drug indicated to treat one disease (source node) is enriched in the differential expression profile of another disease (sink node). Only associations between comorbid diseases ^9^ from different ICD10 categories are shown. The name of the drug connecting both diseases is indicated below the edge. Only associations detected exclusively in one sex are shown, with red (blue) edges denoting associations detected only in females (males). Solid (dashed) lines indicate that the Normalized Enrichment Score of the GSEA is positive (negative), for all other associations see Supplementary Figure 10.

Drug-mediated disease associations may reflect that both diseases share symptoms, patients suffering from one disease also have the other disease, or the treatment of one disease plays a role in the development of the other. In the first scenario, IBS (K58) and major depression (F33), two comorbid diseases with shared pathogenesis ^107^ were connected by lubiprostone, used to treat constipation, a consequence of both diseases ^108^ (see Figure 5). The drug was indicated for treating IBS and enriched in the differential expression profile of major depression in women (but not men). Interestingly, constipation is more likely in women with depression than in men ^109^. In the second one, the association between T2D (E11) and schizophrenia (F20), where drugs used to treat diabetes (metformin and gliclazide) were enriched in women with schizophrenia, could indicate that such patients were indeed taking the aforementioned drugs or suffering from T2D. This hypothesis could be supported by the high representation of elderly patients with schizophrenia in the study dataset, since it has been reported that the prevalence of diabetes in late-life schizophrenia patients is indeed significantly higher in women than in men (35% vs. 21.53%) ^110^. In the third one, it has been described that patients with essential thrombocythemia have an increased risk of transformation to myelofibrosis and that this risk is higher in patients taking anagrelide ^111^ (especially in women), a drug indicated to treat essential thrombocythemia (D47) and enriched in women with myelofibrosis (D75). However, the origin of the data and the lack of metadata preclude further investigation of this hypothesis.

Other comorbidity relationships with pharmacological associations not captured by Westergaard *et al*. ^9^ included the association between T1D and asthma ^112^, where the risk is higher in boys compared to girls ^113^. We found 15 T1D-enriched drugs indicated to treat asthma in men, including salbutamol and salmeterol, not found in women. Interestingly, an increase in blood glucose levels associated with salbutamol use has been described ^114^, highlighting the need to pay special attention to the use of certain treatments depending on the sex of the patients. Similarly, we found two drugs, risperidone and citalopram, significantly enriched in women with schizophrenia, which have also been associated with Alzheimer’s disease. The first is an atypical antipsychotic used to treat schizophrenia ^115^ and behavioral symptoms in Alzheimer’s disease patients ^116^. At the same time, the second is an antidepressant used to treat depression and negative symptoms in schizophrenia ^117^ and agitation in patients with Alzheimer’s disease ^118^. Interestingly, as mentioned, a significant risk of developing Alzheimer’s disease has been described in schizophrenia patients, being higher in women ^41^, and the response to citalopram has also shown sex differences ^119^. These results support the hypothesis that drug-disease associations are different depending on the sex of the patients, and that this may also play a role in comorbidity relationships. However, further studies are needed that directly analyze molecular-level differences in the effect of drugs as a function of the sex of the patient.

### Conclusions

For the first time, disease networks have been generated using separate molecular data for women and men, observing how, based on their differential expression profiles, diseases cluster differently in women and men, being the clusters more heterogeneous in terms of disease categories in women. Calculating transcriptomic similarities between pairs of diseases, we have identified relevant sex-specific differences that align well with those observed at the level of disease co-occurrence. This is the case for T1D and liver cancer in men, or T1D and myocardial infarction and schizophrenia and COPD in women. Interestingly, in the set of comorbidity relationships common to both sexes, we have found differences related to the role of the immune system. Finally, the above-mentioned sex differences have potential consequences for the efficacy of the treatments and the side effects in women and men. The results have shown that it is essential to study comorbidity patterns in women and men separately, investigating the underlying molecular mechanisms and aiming to improve treatments.

## Acknowledgements

J.S.-V. was funded by the Spanish Ministry of Economics and Competitiveness (PID2022-141809OB-I00). J.S.-V. acknowledge funding from the Horizon Europe project COMMUTE under the grant agreement No. 101136957. Funded by the European Union. Views and opinions expressed are however those of the author(s) only and do not necessarily reflect those of the European Union or the European Health and Digital Executive Agency (HADEA). Neither the European Union nor the granting authority can be held responsible for them. L.M.R. was partially funded by the National Institutes of Health (NIH), National Library of Medicine (grants R01-LM011945 and R01-LM012832), by a Fulbright Commission fellowship, and by the National Science Foundation Research Traineeship “Interdisciplinary Training in Complex Networks and Systems” (grant 1735095). J.S.-V., F.X.C., and L.M.R. were funded by the Fundação para a Ciência e a Tecnologia Grant No. 2022.09122.PTDC (https://doi.org/10.54499/2022.09122.PTDC). J.C.-C. was hired under the Generation D initiative, promoted by Red.es, an entity attached to the Ministry for Digital Transformation and the Civil Service, to attract and retain talent through scholarships and training contracts (C005/24-ED CV1), financed by the Recovery, Transformation and Resilience Plan through Next Generation funds. M.F.-R. and R.T.-S. were funded by the Ministry of Education of the Valencian Regional Government (PROMETEO/CIPROM/2022/58). R.T.-S. was funded by the Spanish Ministry of Science, Innovation and Universities (PID2021–129099OB-I00). The funders had no role in study design, data collection and analysis, publication decisions, or manuscript preparation. The authors declare that they have no competing interests. We thank Beatriz Urda (Barcelona Supercomputing Center) for critical reading of the manuscript.

## Data availability

All data needed to understand and assess the conclusions of this research are available in the main text, supplementary materials, and https://github.com/jonsv89/SHDC/blob/main/Datasets.txt. The raw datasets are publicly available and can be downloaded from GEO (https://www.ncbi.nlm.nih.gov/geo) and ArrayExpress (https://www.ebi.ac.uk/arrayexpress).

## Code availability

The code is available at https://github.com/jonsv89/SHDC.

## Author contributions

A.V. and J.S.-V. designed all experiments. J.S.-V., M.F.-R., F.X.C., J.C.-C., and I.N.-C. performed the experiments. D.C., L.M.R., J.C.-C. and R.T-S. provided technical advice. J.S.- V. and AV. wrote the manuscript. All authors discussed the results and commented on the manuscript.

