## Supplementary material for "Sex-specific transcriptome similarity networks elucidate comorbidity relationships"

### Supplementary Note 1

#### Epidemiological networks

The network generated by Hidalgo *et al.* [1] was generated by analyzing data from Medicare, the U.S. government health insurer that has information on 96% of elderly Americans. The records of over 13 million elderly patients (older than 65) from 1990 to 1993 were analyzed. Diagnoses were annotated using ICD9 three-digit codes. Comorbidity relationships were identified by calculating relative risks and phi correlations based on diagnoses conducted on the same day. The resulting network is undirected and consists of 995 ICD9 codes and 291,172 interactions (co-occurrences), with 104,434 of them being significant (diseases co-occurring more than expected by chance, lower confidence interval >1). Unfortunately, we could only access the network generated for the general population, as the networks stratified by sex or race were inaccessible. In contrast, the network generated by Westergaard *et al.* [2] does have comorbidity ratios available for the general population and stratified by sex of patients. In this case, hospital admissions of almost 7 million Danish patients (48% were women) followed for 21 years were analyzed. Diagnoses were noted using 3-digit ICD10 codes. Comorbidity relationships were calculated using relative risks, measuring the risk of diagnosing a disease within five years after diagnosing an index disease. As a result, the generated networks are directed, where A increases the risk of B, but in turn, B may (or may not) increase the risk of A. Since the networks generated using gene expression are not directed, the comorbidity relationships drawn from the Danish population were treated as undirected. A and B will be connected if A (or B) increases the risk of B (or A) significantly. In total, 53,021 co-occurrences were recorded among 1,329 diseases, with 25,707 being significant at the population level (23,708 in women and 22,085 in men, lower confidence interval >1.01).

### Supplementary Note 2

#### Network backbone

Similarity networks are graphs with edges weighted between 0 and 1, where the former denotes a lack of interaction and the latter the strongest possible interaction. Usually, those similarity edge weights vary significantly in a network, which can lead to some being less than indirect interactions [3]. For example, consider that the similarity measure between diseases A and C is 0.1, while between diseases A and B as well as between diseases B and C is 0.2. Consider also that the similarity of a trajectory is given by the minimum edge weight in it; thus, the similarity of A-B-C is 0.2, which is stronger than the direct edge between A and C, which is redundant for recovering the most similar comorbidity trajectories. In this example, we find the ultrametric backbone by keeping only the edges from A-B and B-C [4]. This network backbone is one of the options to sparsify networks in an algebraically principled way based on the identification of edges which obey a generalized triangular inequality in the context of both undirected [3] and directed [5] networks. In the case of the traditional triangular inequality, we recover the metric backbone. In order to identify those backbones, one needs to transform the similarity measures to an isomorphic (not to alter the original network topology) edge distance measure and compute the shortest distance between the node pairs in every edge. If the indirect distance is shorter than the direct distance, the edge is removed until the backbone is found. The edges in a backbone are sufficient for shortest-path computation because they obey a generalized triangular inequality; whereas those removed are considered redundant. Moreover, the backbone has been shown to be the primary transmission subgraph for information diffusion while preserving the community structure [6,7].

Extracting the network backbone for the DTSN, 71% and 69% of the positive interactions were removed in men and women respectively, compared to a reduction of 56% for men and 57% for women in the number of comorbidities in the epidemiology (similar to the one observed in the US population [5]).

This means that with less than half of the comorbidity relations based on transcriptional similarity is necessary to recover the most similar comorbidity trajectories. As a result, the ORs of finding intra- vs inter-category transcriptional similarities over the backbone increased to 4.4 in women and 3.2 in men (the increase in epidemiology was lower, at 2.78 and 2.73 in women and men, respectively). These results denote that there are more redundant edges connecting diseases of different categories than the same in both transcriptional similarity and comorbidity networks, pointing to topological similarities between the two networks.

When calculating the overlap between the epidemiology backbone and the DTSNs of women and men, significant overlaps are still obtained, recovering 38% and 43% of the comorbidities maintained by eliminating redundant edges. This highlights the topological similarity between both networks and the capacity of transcriptional similarity to recover fundamental connections in the flow of information in the comorbidities. Interestingly, around 55% of the backbone comorbidities recovered by the DTSN backbone are shared between women and men, compared to 42% in the original networks(see Supplementary Figure 11).

### Supplementary Note 3

#### Overlap between DTSN and epidemiological networks

Before studying the biological processes potentially involved in the differences in disease co-occurrence between women and men, we evaluated the ability of DTSN to recover the comorbidities described in two distinct populations [1,2]. We found that 55.93% of the diseases co-occurring in the Danish population were transcriptomically similar (see Figure 3C). Similarly, 51.42% of the disease co-occurrence relationships described in 32 million elderly Americans aged 65 or older enrolled in Medicare were identified in the DTSN (see Supplementary Table 1). This result is of great relevance since it is known that the overlap between comorbidity networks is usually low, mainly due to the difference between populations rather than the methods used to generate them [8]. The results obtained are robust as 96.875% (31/32) DTSNs generated using different gene selections and similarity metrics significantly recovered what is described in epidemiology (see Supplementary Table 1). Interestingly, negative connections (representing transcriptional dissimilarities between diseases) did not significantly overlap epidemiological networks (see Supplementary Table 7). The difference between the two epidemiological networks is evident, with the density in the US network being much higher than that of Denmark because the former focuses on patients of an age characterized by a higher prevalence of multimorbidities [9]. Consequently, specific comorbidities are detected only in the US population, as is the case of nervous system diseases (see Supplementary Figure 12). Interestingly, the negative interactions of the generated DTSNs – which we hypothesize connect diseases that co-occur less than expected by chance (known as inverse comorbidity) – did not significantly overlap the epidemiological networks. The overlap between negative interactions and inverse comorbidity relationships (those presenting an RR<1) was not calculated as they are not credible due to the lack of control over the impact of mortality and duration of the study period on detecting inverse comorbidity relationships.

100% of the comorbidities described in both populations between digestive system diseases, skin diseases in Denmark, and infectious and skin diseases in the US are detected in the DTSN (see Supplementary Figure 12). High overlaps are also obtained for neoplasm co-occurrences, the disease category with the highest representation in the DTSN (~30% of nodes), where 75% of co-occurring diseases are found to be significantly similar (34 and 74 comorbidity relationships in Denmark and US). The ICD with the highest number of co-occurrences retrieved by DTSN in the Danish population refers to metastases, followed by nicotine dependence (7), myelodysplastic syndromes (7), and Crohn's disease (6). More than 75% of their co-occurrences are detected. In the case of the US, IBS (42), diseases of the oral soft tissues (41), and COPD (39) are the diseases for which most comorbidities are recovered (56-62%).

### Supplementary Note 4

#### Pathway-based Disease Transcriptomic Similarity Multilayers

The study of transcriptional similarities between diseases on all protein-coding genes may mask the detection of comorbidities that are due to specific pathway categories. To this end, we generated disease networks with the metrics previously used (see methods) for each comparison (sex) using the genes of each Reactome category (29 in total) and the genes associated with mitochondrial processes (extracted from MitoCarta [10]) separately. As a result, pathways-based Disease Transcriptomic Similarity Multilayer Networks were generated for women (wDTSM) and men (mDTSM) separately, where diseases are connected to each other in each layer if the genes involved in the specific category present similar expression alteration patterns. The percentage of interactions between diseases of the same or different categories varies considerably depending on the layer. The gene expression, protein metabolism, cell cycle, and signal transduction layers have the highest density of connections between diseases of the same category in wDTSM and mDTSM (>50%). The signal transduction and immune system layers have the lowest ratio of intra/interconnections (1.69-1.35), while reproduction, drug ADME, and chromatin organization and muscle contraction have the highest ratios (9.89-3.68), meaning that these biological processes tend to connect more diseases from the same category than from different categories (see Supplementary Table 8). When calculating the overlap of each of the individual layers with the epidemiology, we noted that gene expression, the immune system, and protein metabolism recovered more than 70% of the comorbidities recovered when analyzing the complete list of protein-coding genes (see Supplementary Figure 8C-D), pointing to them as the most informative processes in terms of disease co-occurrence. These results agree with those obtained by Dong et al., who described that 51% of the multimorbidities interpretable through SNPs shared at least one HLA-region SNP [11]. In the case of women, reproduction, DNA replication, and drug ADME are the layers that recover comorbidities with a higher ratio of intra- vs. inter-category co-occurrences (13.7,

8.2, and 6.85), while the circadian clock and immune system recover proportionally more similar percentages of intra/inter comorbidities (1.37 and 1.48, see Supplementary Table 8). In the case of women the highest ratios are found in chromatin organization and, as in men, DNA replication and reproduction (2.91, 2.87 and 2.58), with drug ADME and sensory perception showing the lowest ratios (0.97 and 1.03). These results show that 1) the role of different biological processes in comorbidities is different in women and men and 2) that processes such as the immune system, circadian clock, and sensory perception have a greater capacity to recover comorbidities between diseases that affect different systems and therefore could show a more systemic role at the organism level.

Supplementary Figure 13 shows that specific disease pairs, such as ulcerative colitis and IBS, are connected in practically all layers in wDTSM and mDTSM. In contrast, others are only connected in a reduced number of layers (smoking and Crohn's disease). Notably, by generating the different layers, only two (three) comorbidities are lost that are detected when analyzing the complete list of genes in females (males). In contrast, seven new comorbidities are recovered in women and men by generating the different layers, indicated with an asterisk in Supplementary Figure 13 (see also Supplementary Figure 8E-F). The co-occurrence of T1D and pancreatic cancer in women is newly recovered only by the immune system layer, which is known to play a key role in both T1D and cancer [12]. Similarly, the increased risk of vascular disease in women with T1D is newly recovered thanks to the DNA repair layer, a process that has been described to be altered in both diseases [13,14]. In the case of men, the co-occurrence of IgA nephropathy and bladder cancer is recovered by the immune system and cell cycle layers [15,16].

Additionally, the same comorbid diseases can be connected in different layers according to sex. For example, smoking and schizophrenia, for which a bidirectional relationship has been described [17,18], are connected in men in the cell cycle layer. In contrast, in women, they are connected in the layers of the immune system, metabolism, and small molecular transport. Interestingly, previous work has proposed that differences between women and men in the relationship between smoking and schizophrenia may be due to differences in the type of smoking or differences in the immune system, where estrogen tends to increase the immune response whereas testosterone decreases it [19]. Another interesting example is the higher risk of myocardial infarction (I21) in patients with peripheral artery disease (I73) [20]. These two diseases are connected in both sexes in the gene expression transcription layer, and in the hemostasis, signal transduction, and vesicle-mediated transport layers in women, and the immune system, metabolism of proteins and metabolism in men. These processes have been previously related to both diseases and differences between women and men have also been described [21–23]. However, more specific studies are needed to understand the underlying biology better.

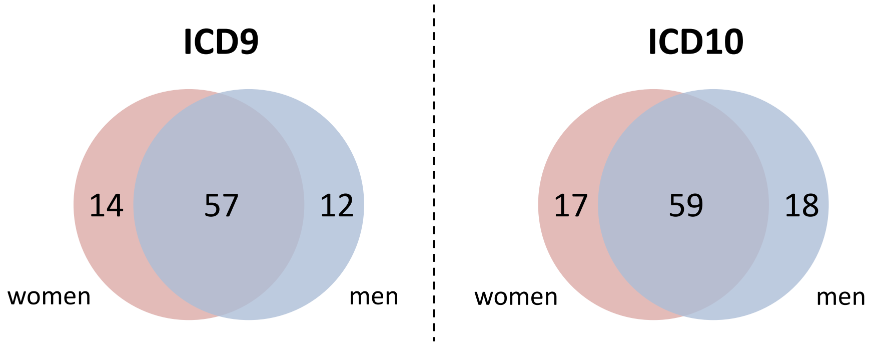


**Supplementary Figure 1. Disease by sex.** Venn diagram showing the number of diseases, coded using versions 9 and 10 of the International Classification of Diseases, with transcriptional information for females (red) and males (blue).

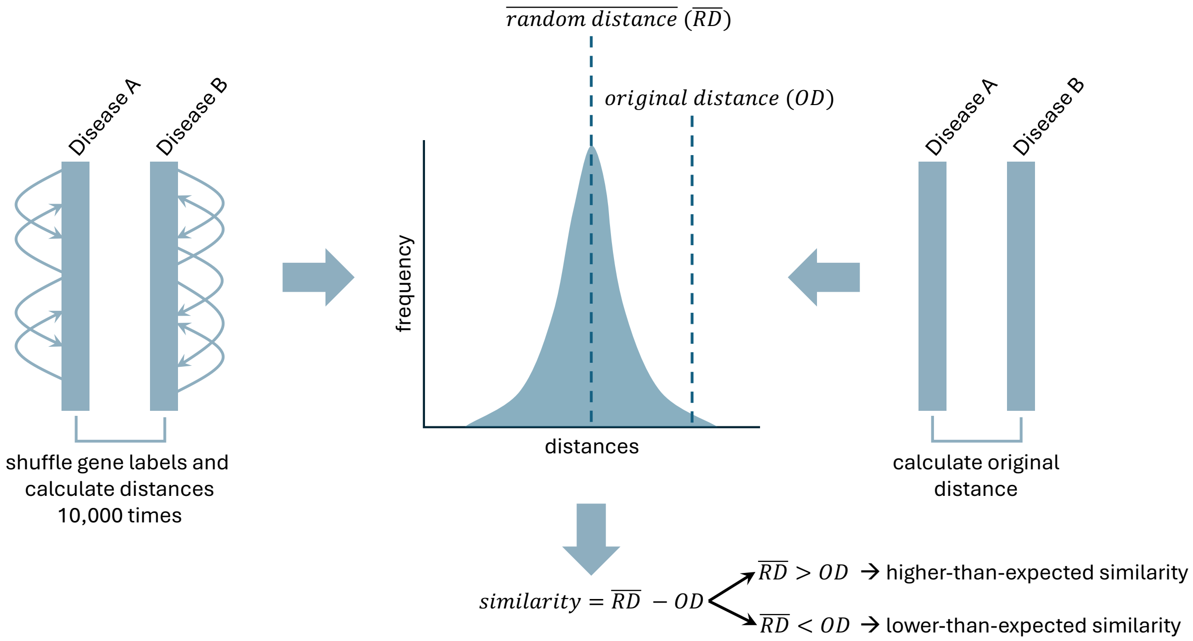


**Supplementary Figure 2. Transformation of distances into similarities**. Euclidean, Canberra, and Manhattan distances were compared with the mean of the random distances (calculated through 10,000 permutations shuffling gene labels), obtaining positive (negative) values.

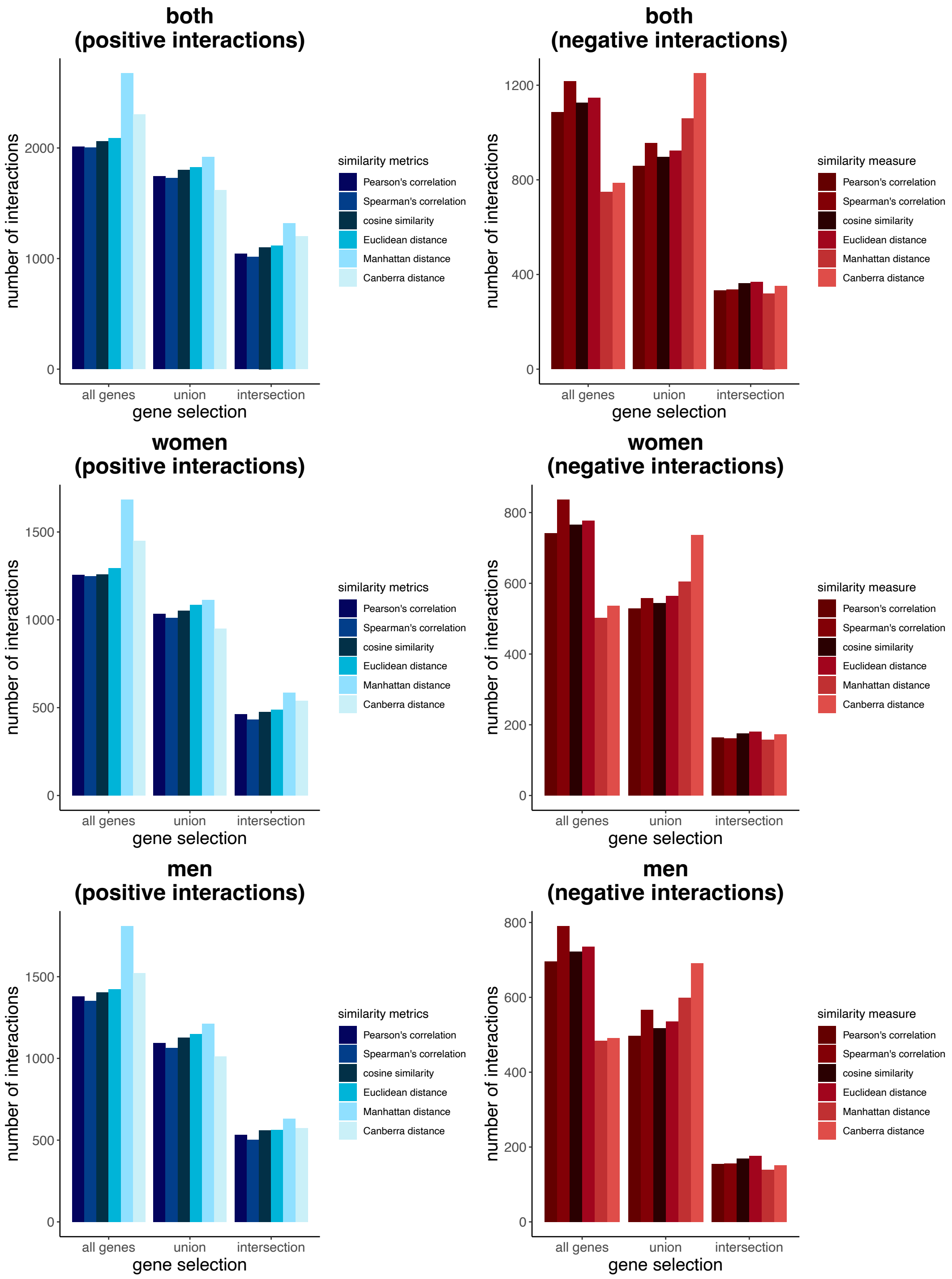


**Supplementary Figure 3.** Number of positive (blue) and negative (red) interactions between diseases when analyzing men and women together (both) or separately.

**Supplementary Table 1.** Recall of the positive interactions in the networks generated using different gene selections (intersection of analyzed genes, union and intersection of sDEGs in each pair of diseases) and metrics (Pearson's and Spearman's correlations, cosine similarity, and Euclidean, Manhattan and Canberra distances) with the epidemiological networks (Hidalgo *et al.*, where diseases are coded using ICD9, and Westergaard *et al.*, where diseases are coded using ICD10) analyzing together women and men (termed both). Additionally, overlaps for women and men are only calculated on the Westergaard *et al.* network as are the only ones providing sex-specific disease co-occurrences. The number of overlapping interactions/comorbidities and the associated p-value are also indicated for each gene selection, metric and epidemiological network. Non-significant overlaps are denoted in grey.

| | | ICD9 | | | ICD10 | | | | | | | | |
|---|---|---|---|---|---|---|---|---|---|---|---|---|---|
| | | both | | | both | | | women | | | men | | |
| | | overlap | recall | p-value | overlap | recall | p-value | overlap | recall | p-value | overlap | recall | p-value |
| **analyzed genes (intersection)** | **Pearson's correlation** | 672 | 50.07 | 0 | 97 | 54.8 | 0.001 | 67 | 52.34 | 0.0126 | 71 | 60.68 | 0 |
| | **Spearman's correlation** | 657 | 48.96 | 3e-04 | 95 | 53.67 | 0.0065 | 65 | 50.78 | 0.0207 | 67 | 57.26 | 3e-04 |
| | **Cosine similarity** | 681 | 50.75 | 0 | 98 | 55.37 | 0.0011 | 66 | 51.56 | 0.0113 | 71 | 60.68 | 0 |
| | **Euclidean distance** | 690 | 51.42 | 0 | 99 | 55.93 | 0.0012 | 68 | 53.12 | 0.0126 | 71 | 60.68 | 0 |
| | **Manhattan distance** | 884 | 65.87 | 9e-04 | 117 | 66.1 | 0.0223 | 80 | 62.5 | 0.2156 | 78 | 66.67 | 0.0175 |
| | **Canberra distance** | 748 | 55.74 | 9e-04 | 104 | 58.76 | 0.0097 | 75 | 58.59 | 0.0261 | 72 | 61.54 | 5e-04 |
| **sDEGs (union)** | **Pearson's correlation** | 594 | 44.26 | 0 | 89 | 50.28 | 0 | 54 | 42.19 | 0.0076 | 56 | 47.86 | 0 |
| | **Spearman's correlation** | 578 | 43.07 | 5e-04 | 87 | 49.15 | 0.0013 | 56 | 43.75 | 0.0014 | 54 | 46.15 | 0 |
| | **Cosine similarity** | 608 | 45.31 | 2e-04 | 88 | 49.72 | 0.0022 | 55 | 42.97 | 0.0093 | 59 | 50.43 | 0 |
| | **Euclidean distance** | 614 | 45.75 | 1e-04 | 91 | 51.41 | 0 | 58 | 45.31 | 0.0044 | 58 | 49.57 | 0 |
| | **Manhattan distance** | 623 | 46.42 | 0.0019 | 94 | 53.11 | 2e-04 | 59 | 46.09 | 0.0026 | 60 | 51.28 | 0 |
| | **Canberra distance** | 534 | 39.79 | 1e-04 | 84 | 47.46 | 0 | 54 | 42.19 | 7e-04 | 50 | 42.74 | 0.0013 |
| **sDEGs (intersection)** | **Pearson's correlation** | 345 | 25.71 | 7e-04 | 69 | 38.98 | 0 | 27 | 21.09 | 0 | 38 | 32.48 | 0 |
| | **Spearman's correlation** | 333 | 24.81 | 0.0046 | 65 | 36.72 | 0 | 25 | 19.53 | 4e-04 | 34 | 29.06 | 0 |
| | **Cosine similarity** | 360 | 26.83 | 2e-04 | 70 | 39.55 | 0 | 27 | 21.09 | 8e-04 | 40 | 34.19 | 0 |
| | **Euclidean distance** | 360 | 26.83 | 6e-04 | 70 | 39.55 | 0 | 28 | 21.88 | 5e-04 | 41 | 35.04 | 0 |
| | **Manhattan distance** | 413 | 30.77 | 0.0256 | 76 | 42.94 | 1e-04 | 30 | 23.44 | 0.0063 | 39 | 33.33 | 0 |
| | **Canberra distance** | 377 | 28.09 | 0.0813 | 74 | 41.81 | 1e-04 | 29 | 22.66 | 0.0017 | 38 | 32.48 | 0 |
| | **Fisher's exact test** | 216 | 16.1 | 2e-04 | 50 | 28.25 | 0 | 21 | 16.41 | 0 | 19 | 16.24 | 0.0027 |

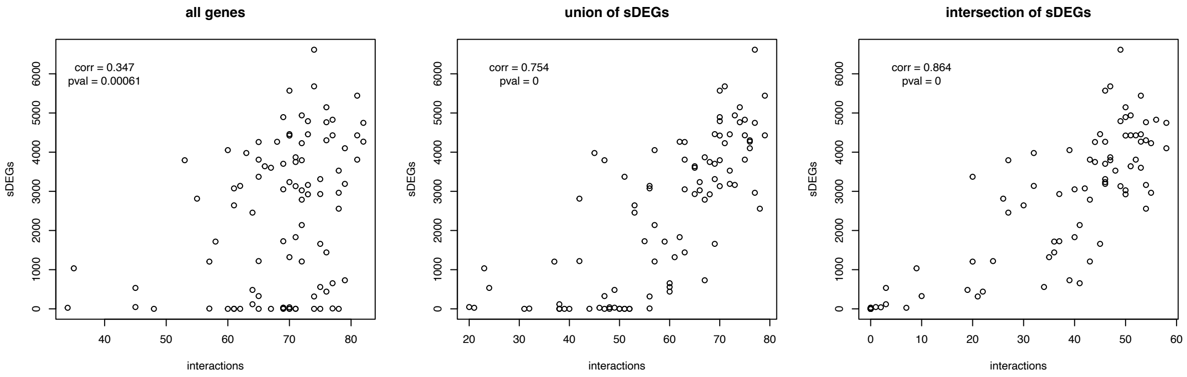


**Supplementary Figure 4.** Correlation between the number of sDEGs and the number of interactions detected when working with the entire list of genes or the union or intersection of the sDEGs.

**Supplementary Table 2.** Odds Ratio of intra- vs. inter-disease category similarities when adjusting for sex differences or analyzing separately men and women.

| | **interactions** | **OR** | **p-value** |
|---|---|---|---|
| Adjusted | positive | 2.21 | $7.93*10^{-19}$ |
| | negative | 0.65 | 0.0001 |
| Women | positive | 2.81 | $3.19*10^{-19}$ |
| | negative | 0.51 | $2.8*10^{-7}$ |
| Men | positive | 1.83 | $1.08*10^{-8}$ |
| | negative | 0.84 | 0.223 |

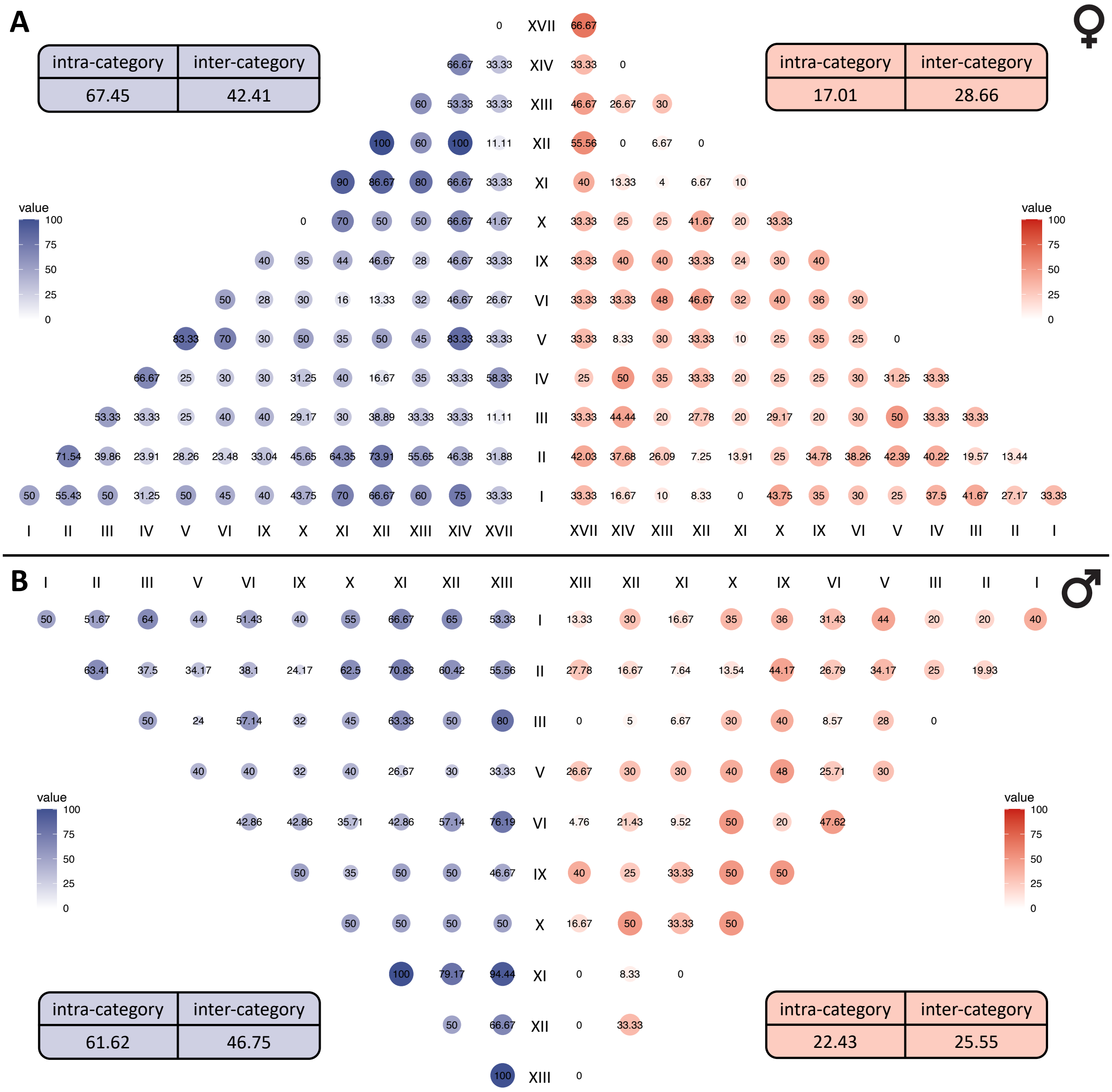


**I** Certain infectious and parasitic diseases
**II** Neoplasms
**III** Diseases of the blood and blood-forming organs and certain disorders involving the immune mechanism
**IV** Endocrine, nutritional and metabolic diseases
**V** Mental and behavioral disorders
**VI** Diseases of the nervous system
**IX** Diseases of the circulatory system
**X** Diseases of the respiratory system
**XI** Diseases of the digestive system
**XII** Diseases of the skin and subcutaneous tissue
**XIII** Diseases of the musculoskeletal system and connective tissue
**XIV** Diseases of the genitourinary system
**XVII** Congenital malformations, deformations and chromosomal abnormalities

**Supplementary Figure 5.** Percentage of intra- and inter-disease category similarities in women (A) and men (B). Positive similarities are denoted in blue and negative similarities in red.

**Supplementary Table 3.** Odds Ratio of finding positive interactions between pairs of disease categories in women compared to men. Significant OR (p-value<=0.05) are highlighted in light blue.

| category 1 | category 2 | OR | p-value |
|---|---|---|---|
| Certain infectious and parasitic diseases | Certain infectious and parasitic diseases | 1.886 | 1 |
| Neoplasms | Certain infectious and parasitic diseases | 1.193 | 0.731 |
| Neoplasms | Neoplasms | 0.868 | 0.69 |
| Diseases of the blood and blood-forming organs | Certain infectious and parasitic diseases | 0.819 | 1 |
| Diseases of the blood and blood-forming organs | Neoplasms | 1.275 | 0.532 |
| Diseases of the blood and blood-forming organs | Diseases of the blood and blood-forming organs | 0.68 | 1 |
| Mental and behavioural disorders | Certain infectious and parasitic diseases | 0.784 | 1 |
| Mental and behavioural disorders | Neoplasms | 0.724 | 0.471 |
| Mental and behavioural disorders | Diseases of the blood and blood-forming organs | 1 | 1 |
| Mental and behavioural disorders | Mental and behavioural disorders | 4.341 | 0.545 |
| Diseases of the nervous system | Certain infectious and parasitic diseases | 0.822 | 1 |
| Diseases of the nervous system | Neoplasms | 0.676 | 0.315 |
| Diseases of the nervous system | Diseases of the blood and blood-forming organs | 0.322 | 0.088 |
| Diseases of the nervous system | Mental and behavioural disorders | 1.538 | 0.741 |
| Diseases of the nervous system | Diseases of the nervous system | 1 | 1 |
| Diseases of the circulatory system | Certain infectious and parasitic diseases | 1.231 | 1 |
| Diseases of the circulatory system | Neoplasms | 1.479 | 0.315 |
| Diseases of the circulatory system | Diseases of the blood and blood-forming organs | 1.652 | 0.561 |
| Diseases of the circulatory system | Mental and behavioural disorders | 0.8 | 1 |
| Diseases of the circulatory system | Diseases of the nervous system | 0.502 | 0.377 |
| Diseases of the circulatory system | Diseases of the circulatory system | 0.68 | 1 |
| Diseases of the respiratory system | Certain infectious and parasitic diseases | 0.615 | 0.724 |
| Diseases of the respiratory system | Neoplasms | 0.619 | 0.225 |
| Diseases of the respiratory system | Diseases of the blood and blood-forming organs | 0.532 | 0.514 |
| Diseases of the respiratory system | Mental and behavioural disorders | 1.276 | 1 |
| Diseases of the respiratory system | Diseases of the nervous system | 0.8 | 1 |
| Diseases of the respiratory system | Diseases of the circulatory system | 1 | 1 |
| Diseases of the respiratory system | Diseases of the respiratory system | 0 | 0.182 |
| Diseases of the digestive system | Certain infectious and parasitic diseases | 1.249 | 1 |
| Diseases of the digestive system | Neoplasms | 0.635 | 0.239 |
| Diseases of the digestive system | Diseases of the blood and blood-forming organs | 0.272 | 0.046 |
| Diseases of the digestive system | Mental and behavioural disorders | 1.249 | 1 |
| Diseases of the digestive system | Diseases of the nervous system | 0.213 | 0.032 |
| Diseases of the digestive system | Diseases of the circulatory system | 0.623 | 0.572 |
| Diseases of the digestive system | Diseases of the respiratory system | 3.385 | 0.111 |
| Diseases of the digestive system | Diseases of the digestive system | 0 | 1 |

**Supplementary Table 4.** Odds Ratio of finding negative interactions between pairs of disease categories in women compared to men. Significant OR (p-value<=0.05) are highlighted in light blue.

| category 1 | category 2 | OR | p-value |
|---|---|---|---|
| Certain infectious and parasitic diseases | Certain infectious and parasitic diseases | 0.53 | 1 |
| Neoplasms | Certain infectious and parasitic diseases | 1.518 | 0.414 |
| Neoplasms | Neoplasms | 0.904 | 0.874 |
| Diseases of the blood and blood-forming organs | Certain infectious and parasitic diseases | 1.596 | 0.731 |
| Diseases of the blood and blood-forming organs | Neoplasms | 0.646 | 0.29 |
| Diseases of the blood and blood-forming organs | Diseases of the blood and blood-forming organs | Inf | 0.087 |
| Mental and behavioural disorders | Certain infectious and parasitic diseases | 0.566 | 0.704 |
| Mental and behavioural disorders | Neoplasms | 1.797 | 0.149 |
| Mental and behavioural disorders | Diseases of the blood and blood-forming organs | 1.965 | 0.501 |
| Mental and behavioural disorders | Mental and behavioural disorders | 0 | 1 |
| Diseases of the nervous system | Certain infectious and parasitic diseases | 1 | 1 |
| Diseases of the nervous system | Neoplasms | 1.368 | 0.419 |
| Diseases of the nervous system | Diseases of the blood and blood-forming organs | 3.367 | 0.171 |
| Diseases of the nervous system | Mental and behavioural disorders | 1.859 | 0.695 |
| Diseases of the nervous system | Diseases of the nervous system | 0.657 | 1 |
| Diseases of the circulatory system | Certain infectious and parasitic diseases | 1 | 1 |
| Diseases of the circulatory system | Neoplasms | 0.56 | 0.087 |
| Diseases of the circulatory system | Diseases of the blood and blood-forming organs | 0.481 | 0.364 |
| Diseases of the circulatory system | Mental and behavioural disorders | 0.665 | 0.748 |
| Diseases of the circulatory system | Diseases of the nervous system | 2.213 | 0.345 |
| Diseases of the circulatory system | Diseases of the circulatory system | 0.68 | 1 |
| Diseases of the respiratory system | Certain infectious and parasitic diseases | 1.286 | 1 |
| Diseases of the respiratory system | Neoplasms | 2.007 | 0.183 |
| Diseases of the respiratory system | Diseases of the blood and blood-forming organs | 0.591 | 0.716 |
| Diseases of the respiratory system | Mental and behavioural disorders | 0.74 | 1 |
| Diseases of the respiratory system | Diseases of the nervous system | 0.554 | 0.527 |
| Diseases of the respiratory system | Diseases of the circulatory system | 0.438 | 0.333 |
| Diseases of the respiratory system | Diseases of the respiratory system | 0.53 | 1 |
| Diseases of the digestive system | Certain infectious and parasitic diseases | 0 | 0.231 |
| Diseases of the digestive system | Neoplasms | 2.004 | 0.233 |
| Diseases of the digestive system | Diseases of the blood and blood-forming organs | 5.812 | 0.189 |
| Diseases of the digestive system | Mental and behavioural disorders | 0.637 | 1 |
| Diseases of the digestive system | Diseases of the nervous system | 10.814 | 0.023 |
| Diseases of the digestive system | Diseases of the circulatory system | 0.815 | 1 |
| Diseases of the digestive system | Diseases of the respiratory system | 0.384 | 0.301 |
| Diseases of the digestive system | Diseases of the digestive system | Inf | 1 |

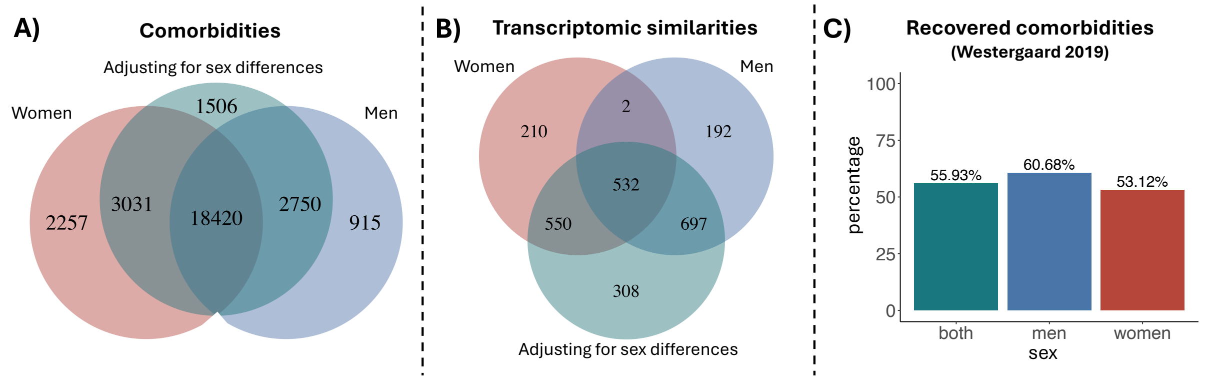


**Supplementary Figure 6.** A-B) Venn diagrams indicating the number of comorbidities [2] (A) and significant disease transcriptomic similarities (B) shared when performing the analyses separately for women (red) and men (blue) and adjusting for sex differences (green). C) Barplots indicating the percentage of comorbidities [2] recovered by calculating transcriptomic similarities between diseases separately for women (red) and men (blue) and adjusting for sex differences (green).

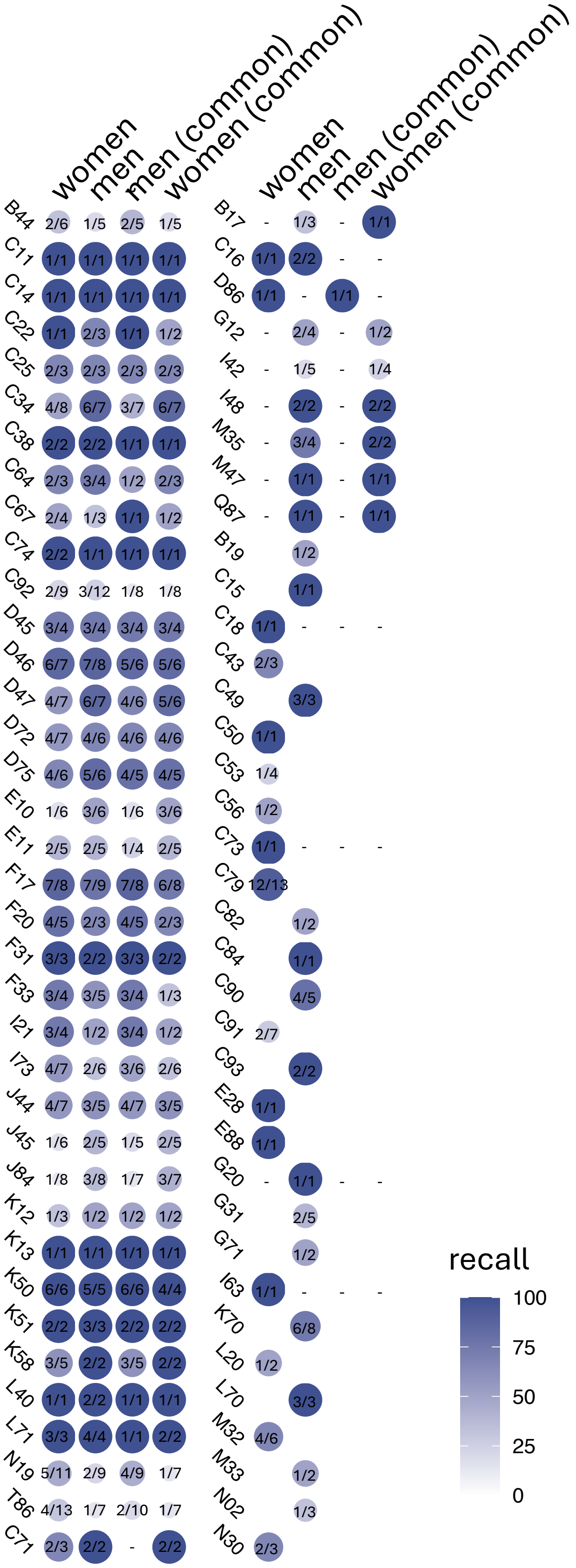

**Supplementary Figure 7.** Recall of comorbidities described for each disease separately for women and men (first two columns), considering also those diseases with information for both sexes (last two columns). The size and color of the circle indicate the recall obtained. Within each circle, the number of comorbidities recovered from the total described for each disease in each sex is shown. The "-" indicates that no comorbidities have been described for that disease in the corresponding sex.

**Supplementary Table 5.** Number of comorbidities (and the associated recall, precision and p-value) recovered by the networks generated for men and women focused on the genes associated to different Reactome categories.

| | women | | | | men | | | |
|---|---|---|---|---|---|---|---|---|
| | number | recall | p-value | precision | number | recall | p-value | precision |
| Drug ADME | 6 | 4.69 | 7e-04 | 30 | 7 | 5.98 | 0 | 30.43 |
| Circadian Clock | 10 | 7.81 | 0 | 14.49 | 15 | 12.82 | 0 | 14.56 |
| Cell-Cell communication | 10 | 7.81 | 2e-04 | 16.39 | 14 | 11.97 | 0 | 15.73 |
| Reproduction | 11 | 8.59 | 0 | 16.18 | 12 | 10.26 | 0 | 10.34 |
| Autophagy | 14 | 10.94 | 0 | 14.43 | 11 | 9.4 | 2e-04 | 9.82 |
| Neuronal System | 17 | 13.28 | 0 | 12.88 | 22 | 18.8 | 0 | 13.58 |
| Muscle contraction | 18 | 14.06 | 0 | 27.69 | 15 | 12.82 | 0 | 17.65 |
| Sensory Perception | 18 | 14.06 | 0 | 21.95 | 18 | 15.38 | 0 | 18.18 |
| Organelle biogenesis and maintenance | 21 | 16.41 | 0.0011 | 7.69 | 25 | 21.37 | 0 | 7.74 |
| Programmed Cell Death | 22 | 17.19 | 0 | 8.87 | 23 | 19.66 | 0 | 7.96 |
| Protein localization | 24 | 18.75 | 0 | 11.71 | 19 | 16.24 | 0 | 9.6 |
| Chromatin organization | 27 | 21.09 | 0 | 12.05 | 26 | 22.22 | 0 | 10.57 |
| Extracellular matrix organization | 27 | 21.09 | 0 | 9.12 | 28 | 23.93 | 0 | 8.7 |
| DNA_Replication | 28 | 21.88 | 0 | 8.07 | 29 | 24.79 | 0 | 7.29 |
| Hemostasis | 30 | 23.44 | 3e-04 | 7.44 | 36 | 30.77 | 0 | 7.33 |
| Transport of small molecules | 31 | 24.22 | 0 | 8.54 | 31 | 26.5 | 0 | 8.49 |
| Vesicle-mediated transport | 34 | 26.56 | 0 | 7.14 | 39 | 33.33 | 0 | 6.44 |
| Developmental Biology | 34 | 26.56 | 0 | 7.41 | 37 | 31.62 | 0 | 6.85 |
| DNA Repair | 35 | 27.34 | 0 | 7.17 | 36 | 30.77 | 0 | 6.5 |
| Metabolism of RNA | 38 | 29.69 | 0.0052 | 6.51 | 41 | 35.04 | 0 | 6.21 |
| Cellular responses to stimuli | 39 | 30.47 | 1e-04 | 7.08 | 48 | 41.03 | 0 | 7 |
| Cell Cycle | 45 | 35.16 | 5e-04 | 6.06 | 43 | 36.75 | 1e-04 | 4.85 |
| Mitochondria All | 47 | 36.72 | 0 | 6.76 | 45 | 38.46 | 0 | 6.44 |
| Metabolism of proteins | 48 | 37.5 | 0.0018 | 5.9 | 52 | 44.44 | 0 | 5.76 |
| Disease | 48 | 37.5 | 0 | 6.51 | 44 | 37.61 | 0 | 5.33 |
| Signal Transduction | 49 | 38.28 | 8e-04 | 5.86 | 49 | 41.88 | 0 | 5.11 |
| Metabolism | 50 | 39.06 | 0 | 6.84 | 47 | 40.17 | 0 | 5.77 |
| Immune System | 52 | 40.62 | 3e-04 | 5.79 | 53 | 45.3 | 0 | 5.23 |
| Gene expression Transcription | 53 | 41.41 | 0 | 6.49 | 55 | 47.01 | 0 | 6.17 |

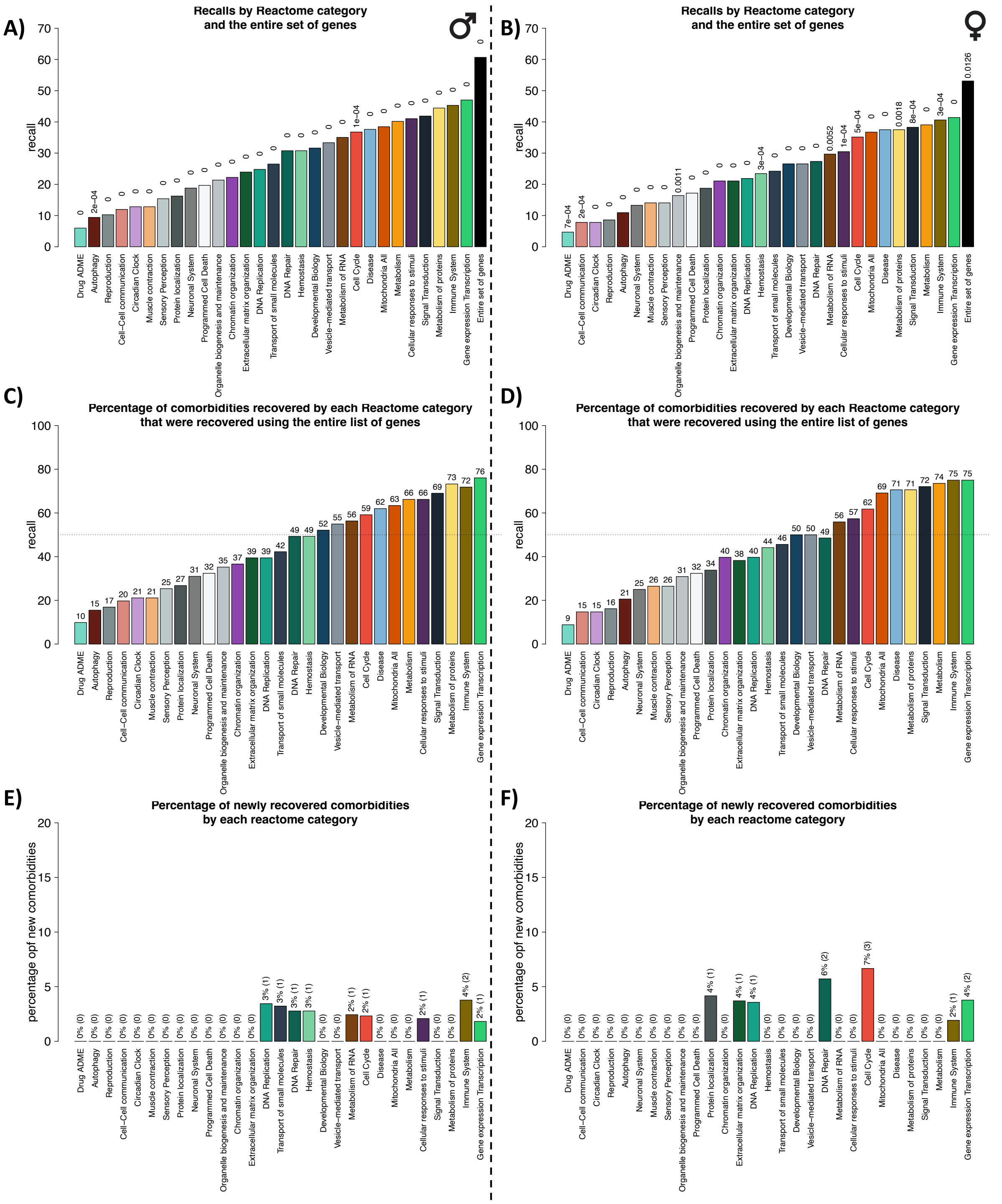


**Supplementary Figure 8. Comorbidities recovered by analyzing mitochondria-associated and reactome category genes.** A and B) Percentage of comorbidities described by Westergaard *et al*. recovered (recall) by calculating the similarities between diseases using the genes of each Reactome category separately, the genes associated with mitochondrial processes (mitochondrial), or the complete list of genes in men (A) and women (B), indicating the pvalue of the overlap with the epidemiology. C and D) Percentage of comorbidities recovered using the complete list of genes that are recovered using only genes from each Reactome category in men (C) and women (D). C and D) Percentage of new comorbidities that are recovered by analyzing the similarities between diseases using only the genes in each Reactome category and that were not recovered using the complete list of genes in men (E) and women (F).

**Supplementary Table 6.** Jaccard index (JI) of the Reactome pathways enriched (up or down) in women and men by disease. The number of pathways up- and down-regulated in each disease in men (women) is indicated separated by a "-".

| | JI up | JI down | Nº up (men-women) | Nº down (men-women) |
|---|---|---|---|---|
| C11 | 0.72 (169/234) | 0.33 (5/15) | 213-190 | 10-10 |
| C14 | 0.68 (190/280) | 0.62 (25/40) | 234-236 | 38-27 |
| C25 | 0.78 (389/496) | 0.58 (7/12) | 471-414 | 10-9 |
| C34 | 0.8 (207/260) | 0.28 (51/181) | 255-212 | 54-178 |
| C38 | 0.07 (6/87) | 0.11 (2/19) | 84-9 | 18-3 |
| C64 | 0.8 (287/357) | 0.81 (44/54) | 297-347 | 45-53 |
| C67 | 0.01 (2/154) | 0.21 (6/29) | 154-2 | 27-8 |
| C74 | 0.57 (100/175) | 0.17 (11/63) | 165-110 | 38-36 |
| C92 | 0.28 (41/148) | 0.15 (10/65) | 142-47 | 12-63 |
| D45 | 0.14 (1/7) | 0.3 (31/105) | 7-1 | 96-40 |
| D46 | 0.01 (1/89) | 0.02 (1/43) | 1-89 | 41-3 |
| D47 | 0.47 (9/19) | 0.66 (264/397) | 16-12 | 304-357 |
| D72 | 0 (0/70) | 0 (0/16) | 2-68 | 14-2 |
| D75 | 0.01 (1/67) | 0.21 (22/103) | 1-67 | 33-92 |
| E11 | 0.03 (1/36) | 0 (0/9) | 14-23 | 7-2 |
| F17 | 0.39 (9/23) | 0.24 (35/143) | 13-19 | 50-128 |
| F20 | 0.21 (18/85) | 0.53 (134/252) | 22-81 | 199-187 |
| F31 | 0 (0/1) | 0 (0/126) | 0-1 | 0-126 |
| F33 | - (0/0) | 0 (0/117) | 0-0 | 0-117 |
| I21 | 0.02 (1/44) | - (0/0) | 44-1 | 0-0 |
| I73 | 0.05 (6/117) | 0 (0/2) | 102-21 | 2-0 |
| J44 | 0.06 (1/17) | 0.13 (6/45) | 4-14 | 11-40 |
| J45 | 0 (0/39) | 0.3 (23/76) | 39-0 | 74-25 |
| K12 | 0.27 (50/186) | 0 (0/8) | 156-80 | 2-6 |
| K13 | 0.34 (31/90) | 0.41 (15/37) | 83-38 | 21-31 |
| K50 | 0.39 (61/156) | 0.69 (11/16) | 136-81 | 13-14 |
| K51 | 0.41 (90/219) | 0.32 (9/28) | 94-215 | 19-18 |
| K58 | 0.57 (137/240) | 0.61 (20/33) | 187-190 | 30-23 |
| L71 | 0.56 (91/162) | 0 (0/29) | 125-128 | 0-29 |
| N19 | 0.29 (6/21) | - (0/0) | 19-8 | 0-0 |

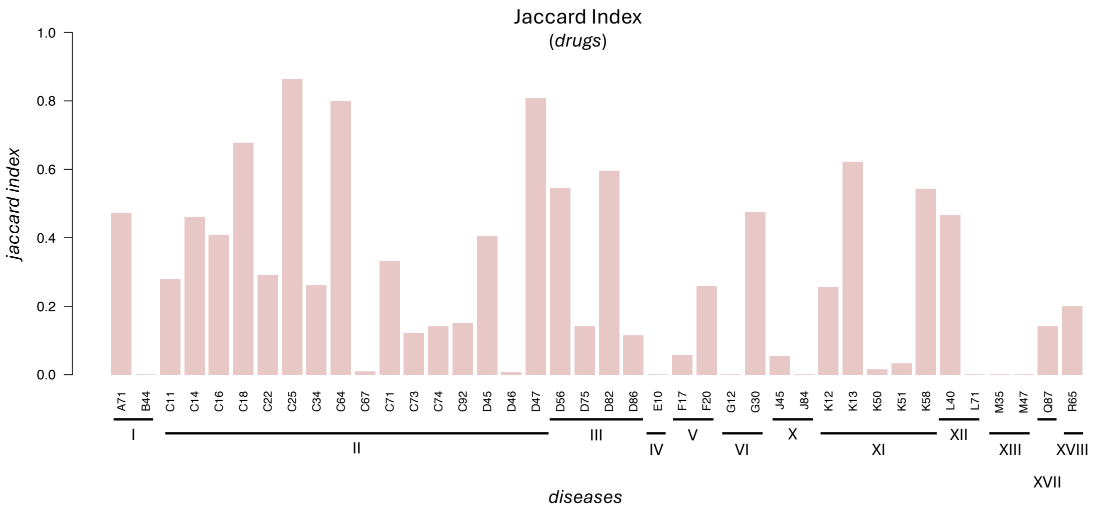


**Supplementary Figure 9**. Jaccard indexes of the drugs associated with women and men in each disease (bar). Horizontal lines below ICD10 codes group the diseases that belong to the same category.

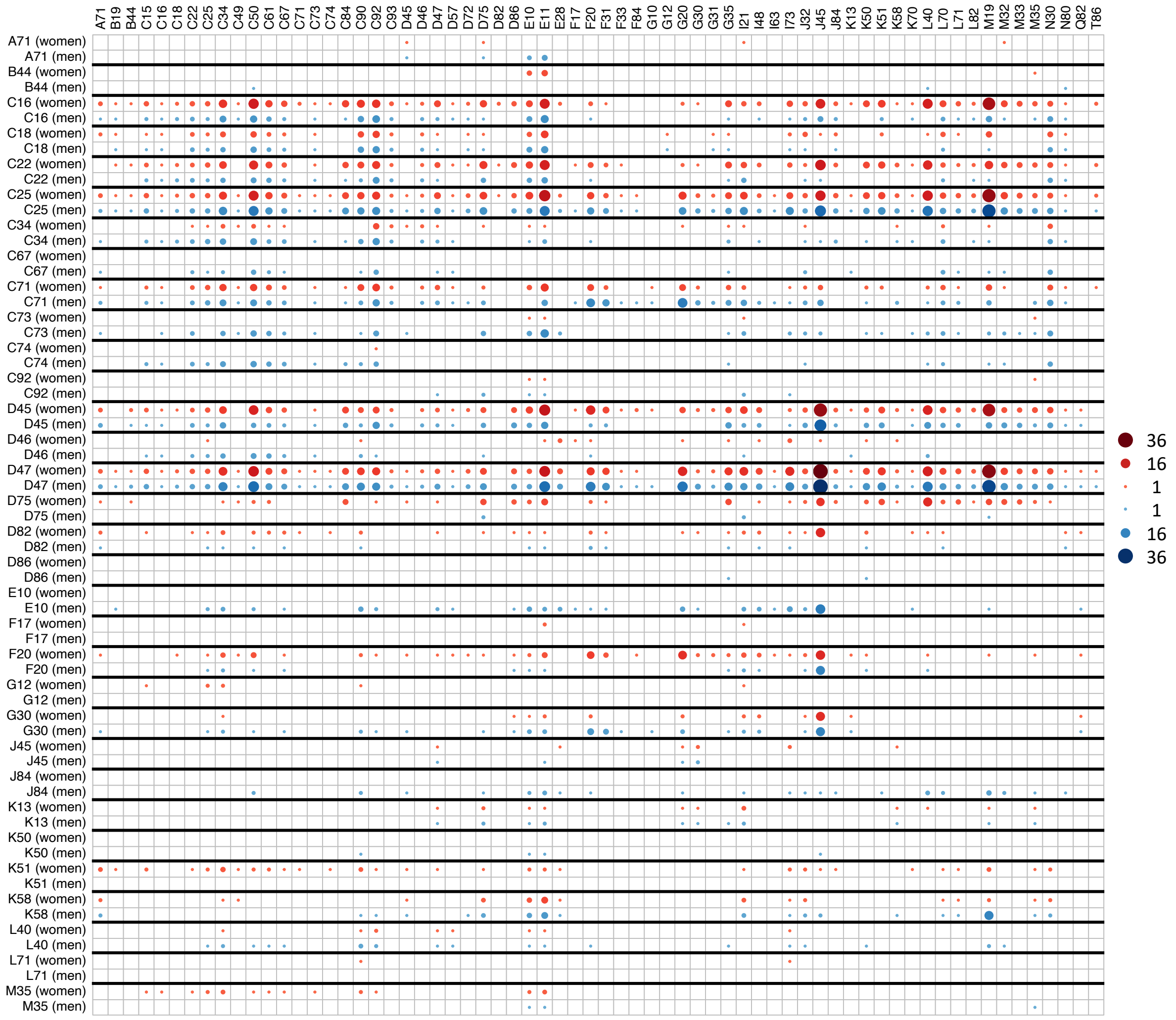

**Supplementary Figure 10. Associations between diseases identified by pharmacological relationships.** Heatmap representation of the relationship between diseases as a function of shared drugs found to be enriched in genes differentially expressed in women and men (rows) and indicated to treat them (columns, extracted from SIDER). The size of the circles denotes the number of drugs found in the association between diseases in women (red) and men (blue).

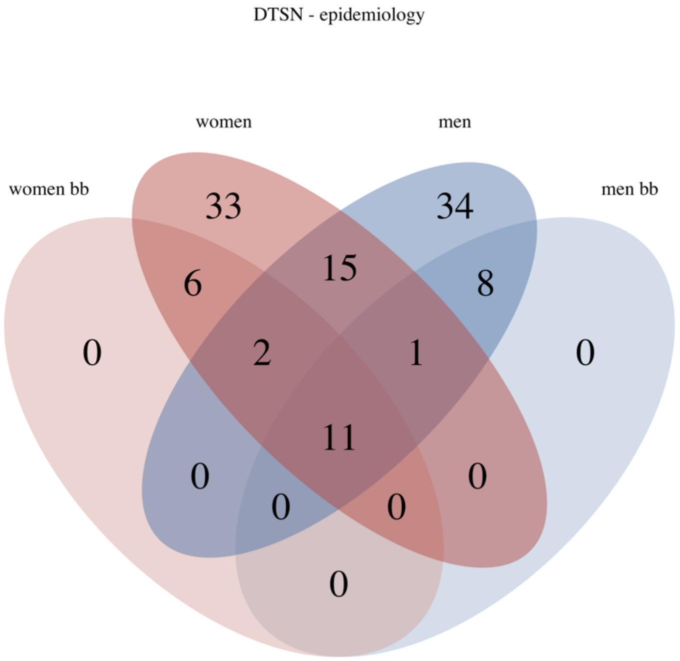


**Supplementary Figure 11**. Venn diagram showing the overlap between DTSNs and epidemiology in women (red) and men (blue), both when analyzing the whole network and the backbone of both networks (women bb and men bb).

**Supplementary Table 7.** Recall of the negative interactions in the networks generated using different gene selections (intersection of analyzed genes, union and intersection of sDEGs in each pair of diseases) and metrics (Pearson's and Spearman's correlations, cosine similarity, and Euclidean, Manhattan and Canberra distances) with the epidemiological networks (Hidalgo *et al.*, where diseases are coded using ICD9, and Westergaard *et al.*, where diseases are coded using ICD10) analyzing together women and men (termed both). Additionally, overlaps for women and men are only calculated on the Westergaard *et al.* network as are the only ones providing sex-specific disease co-occurrences. The number of overlapping interactions/comorbidities and the associated p-value are also indicated for each gene selection, metric and epidemiological network.

| | | ICD9 | | | ICD10 | | | | | | | | |
|---|---|---|---|---|---|---|---|---|---|---|---|---|---|
| | | both | | | both | | | women | | | men | | |
| | | **overlap** | **recall** | **p-value** | **overlap** | **recall** | **p-value** | **overlap** | **recall** | **p-value** | **overlap** | **recall** | **p-value** |
| **analyzed genes (intersection)** | **Pearson's correlation** | 305 | 22.73 | 0.9052 | 44 | 24.86 | 0.8165 | 40 | 31.25 | 0.259 | 27 | 23.08 | 0.94 |
| | **Spearman's correlation** | 334 | 24.89 | 0.9549 | 49 | 27.68 | 0.8086 | 43 | 33.59 | 0.2835 | 32 | 27.35 | 0.8739 |
| | **Cosine similarity** | 317 | 23.62 | 0.965 | 44 | 24.86 | 0.889 | 40 | 31.25 | 0.3359 | 30 | 25.64 | 0.8843 |
| | **Euclidean distance** | 319 | 23.77 | 0.9648 | 44 | 24.86 | 0.9082 | 41 | 32.03 | 0.2998 | 31 | 26.5 | 0.8529 |
| | **Manhattan distance** | 218 | 16.24 | 0.6106 | 33 | 18.64 | 0.594 | 27 | 21.09 | 0.4725 | 23 | 19.66 | 0.5275 |
| | **Canberra distance** | 224 | 16.69 | 0.8969 | 36 | 20.34 | 0.5535 | 31 | 24.22 | 0.1847 | 26 | 22.22 | 0.2627 |
| **sDEGs (union)** | **Pearson's correlation** | 239 | 17.81 | 0.8585 | 36 | 20.34 | 0.6347 | 24 | 18.75 | 0.5653 | 21 | 17.95 | 0.7987 |
| | **Spearman's correlation** | 260 | 19.37 | 0.9196 | 42 | 23.73 | 0.4388 | 23 | 17.97 | 0.7075 | 22 | 18.8 | 0.8805 |
| | **Cosine similarity** | 250 | 18.63 | 0.914 | 38 | 21.47 | 0.6186 | 24 | 18.75 | 0.577 | 20 | 17.09 | 0.9018 |
| | **Euclidean distance** | 255 | 19 | 0.9363 | 39 | 22.03 | 0.6504 | 25 | 19.53 | 0.5423 | 21 | 17.95 | 0.9018 |
| | **Manhattan distance** | 271 | 20.19 | 0.9974 | 42 | 23.73 | 0.8471 | 27 | 21.09 | 0.6663 | 20 | 17.09 | 0.9835 |
| | **Canberra distance** | 338 | 25.19 | 0.9973 | 45 | 25.42 | 0.9578 | 30 | 23.44 | 0.7784 | 21 | 17.95 | 0.9905 |
| **sDEGs (intersection)** | **Pearson's correlation** | 110 | 8.2 | 0.6964 | 13 | 7.34 | 0.8458 | 6 | 4.69 | 0.7728 | 6 | 5.13 | 0.855 |
| | **Spearman's correlation** | 104 | 7.75 | 0.7121 | 14 | 7.91 | 0.7475 | 6 | 4.69 | 0.7355 | 5 | 4.27 | 0.9493 |
| | **Cosine similarity** | 116 | 8.64 | 0.7275 | 16 | 9.04 | 0.6749 | 6 | 4.69 | 0.8361 | 6 | 5.13 | 0.9094 |
| | **Euclidean distance** | 123 | 9.17 | 0.5916 | 16 | 9.04 | 0.715 | 6 | 4.69 | 0.8294 | 7 | 5.98 | 0.8727 |
| | **Manhattan distance** | 96 | 7.15 | 0.5146 | 16 | 9.04 | 0.566 | 6 | 4.69 | 0.7744 | 6 | 5.13 | 0.7713 |
| | **Canberra distance** | 104 | 7.75 | 0.7182 | 17 | 9.6 | 0.5001 | 6 | 4.69 | 0.8199 | 7 | 5.98 | 0.6004 |
| | **Fisher's exact test** | 58 | 4.32 | 0.6793 | 5 | 2.82 | 0.9232 | 3 | 2.34 | 0.7161 | 1 | 0.85 | 0.923 |

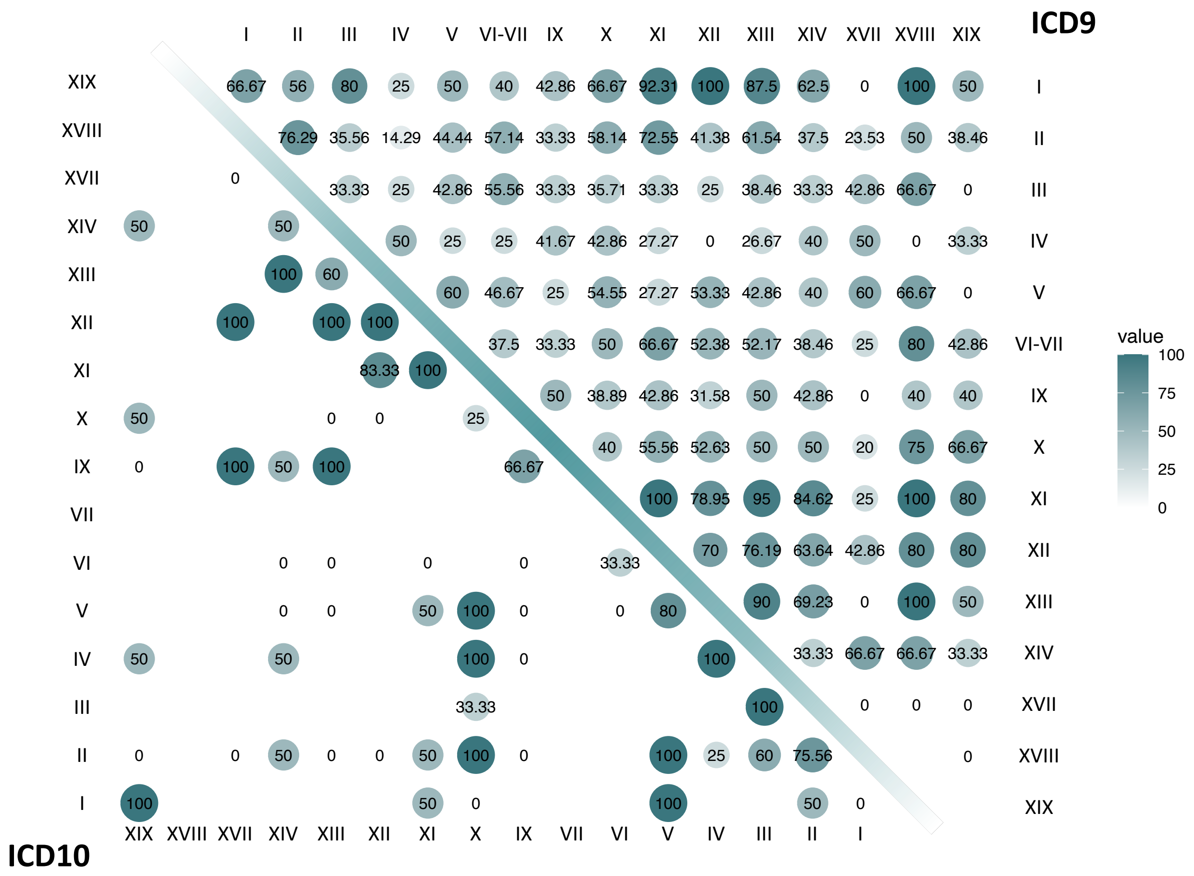


**I** Certain infectious and parasitic diseases
**II** Neoplasms
**III** Diseases of the blood and blood-forming organs and certain disorders involving the immune mechanism
**IV** Endocrine, nutritional and metabolic diseases
**V** Mental and behavioral disorders
**VI** Diseases of the nervous system
**VII** Diseases of the eye and adnexa
**IX** Diseases of the circulatory system
**X** Diseases of the respiratory system
**XI** Diseases of the digestive system
**XII** Diseases of the skin and subcutaneous tissue
**XIII** Diseases of the musculoskeletal system and connective tissue
**XIV** Diseases of the genitourinary system
**XVII** Congenital malformations, deformations and chromosomal abnormalities
**XVIII** Symptoms, signs and abnormal clinical and laboratory findings
**XIX** Injury and poisoning

**Supplementary Figure 12.** Recall of the DTSN (generated adjusting for sex-differences) on the Denmark population (ICD10) and US elderly population (ICD9).

**Supplementary Table 8.** Percentage of intra- and inter-disease category transcriptomic similarities and recovered comorbidity by the different Reactome categories and pathways associated with mitochondria.

| | **similarities** | | | | **recovered comorbidities** | | | |
|---|---|---|---|---|---|---|---|---|
| | **women** | | **men** | | **women** | | **men** | |
| | **intra** | **inter** | **intra** | **inter** | **intra** | **inter** | **intra** | **inter** |
| Autophagy | 9.38 | 2.59 | 7.3 | 3.33 | 14.81 | 8.11 | 11.76 | 7.58 |
| Cell Cycle | 51.03 | 22.68 | 55.14 | 26.72 | 55.56 | 20.27 | 49.02 | 27.27 |
| Cell-Cell communication | 5.28 | 1.71 | 6.76 | 2.5 | 11.11 | 5.41 | 15.69 | 9.09 |
| Cellular responses to stimuli | 43.99 | 15.98 | 42.16 | 20.74 | 44.44 | 20.27 | 50.98 | 33.33 |
| Chromatin organization | 25.81 | 5.42 | 26.22 | 5.83 | 37.04 | 9.46 | 35.29 | 12.12 |
| Circadian Clock | 5.87 | 1.95 | 6.76 | 3.05 | 9.26 | 6.76 | 15.69 | 10.61 |
| Developmental Biology | 33.14 | 13.79 | 32.97 | 16.35 | 38.89 | 17.57 | 41.18 | 24.24 |
| Disease | 48.39 | 22.8 | 41.62 | 26.29 | 51.85 | 27.03 | 52.94 | 25.76 |
| DNA Repair | 42.23 | 13.71 | 44.59 | 15.22 | 44.44 | 14.86 | 43.14 | 21.21 |
| DNA Replication | 37.24 | 8.77 | 36.49 | 10.29 | 44.44 | 5.41 | 39.22 | 13.64 |
| Drug ADME | 2.64 | 0.44 | 2.16 | 0.59 | 9.26 | 1.35 | 5.88 | 6.06 |
| Extracellular matrix organization | 26.98 | 8.13 | 28.65 | 8.45 | 37.04 | 9.46 | 33.33 | 16.67 |
| Gene expression Transcription | 55.13 | 25.07 | 51.89 | 27.35 | 59.26 | 28.38 | 60.78 | 36.36 |
| Hemostasis | 34.02 | 11.44 | 32.7 | 14.48 | 38.89 | 12.16 | 39.22 | 24.24 |
| Immune System | 49.27 | 29.1 | 44.86 | 33.14 | 50 | 33.78 | 54.9 | 37.88 |
| Metabolism | 47.8 | 22.64 | 45.14 | 25.31 | 53.7 | 28.38 | 54.9 | 28.79 |
| Metabolism of proteins | 51.91 | 25.39 | 50 | 28.09 | 50 | 28.38 | 60.78 | 31.82 |
| Metabolism of RNA | 46.92 | 16.9 | 41.62 | 19.8 | 46.3 | 17.57 | 50.98 | 22.73 |
| Mitochondria All | 47.21 | 21.28 | 42.97 | 21.13 | 51.85 | 25.68 | 52.94 | 27.27 |
| Muscle contraction | 6.74 | 1.67 | 8.38 | 2.11 | 20.37 | 9.46 | 17.65 | 9.09 |
| Neuronal System | 13.2 | 3.47 | 13.78 | 4.34 | 18.52 | 9.46 | 27.45 | 12.12 |
| Organelle biogenesis and maintenance | 26.39 | 7.29 | 28.11 | 8.57 | 25.93 | 9.46 | 29.41 | 15.15 |
| Programmed Cell Death | 19.65 | 7.21 | 17.84 | 8.72 | 25.93 | 10.81 | 27.45 | 13.64 |
| Protein localization | 22.29 | 5.14 | 17.03 | 5.28 | 33.33 | 8.11 | 21.57 | 12.12 |
| Reproduction | 11.44 | 1.16 | 15.41 | 2.31 | 18.52 | 1.35 | 15.69 | 6.06 |
| Sensory Perception | 8.8 | 2.07 | 5.95 | 3.01 | 20.37 | 9.46 | 15.69 | 15.15 |
| Signal Transduction | 51.91 | 26.27 | 47.3 | 30.63 | 53.7 | 27.03 | 52.94 | 33.33 |
| Transport of small molecules | 25.51 | 11 | 22.16 | 11.07 | 33.33 | 17.57 | 31.37 | 22.73 |
| Vesicle-mediated transport | 38.12 | 13.79 | 38.38 | 18.15 | 38.89 | 17.57 | 45.1 | 24.24 |

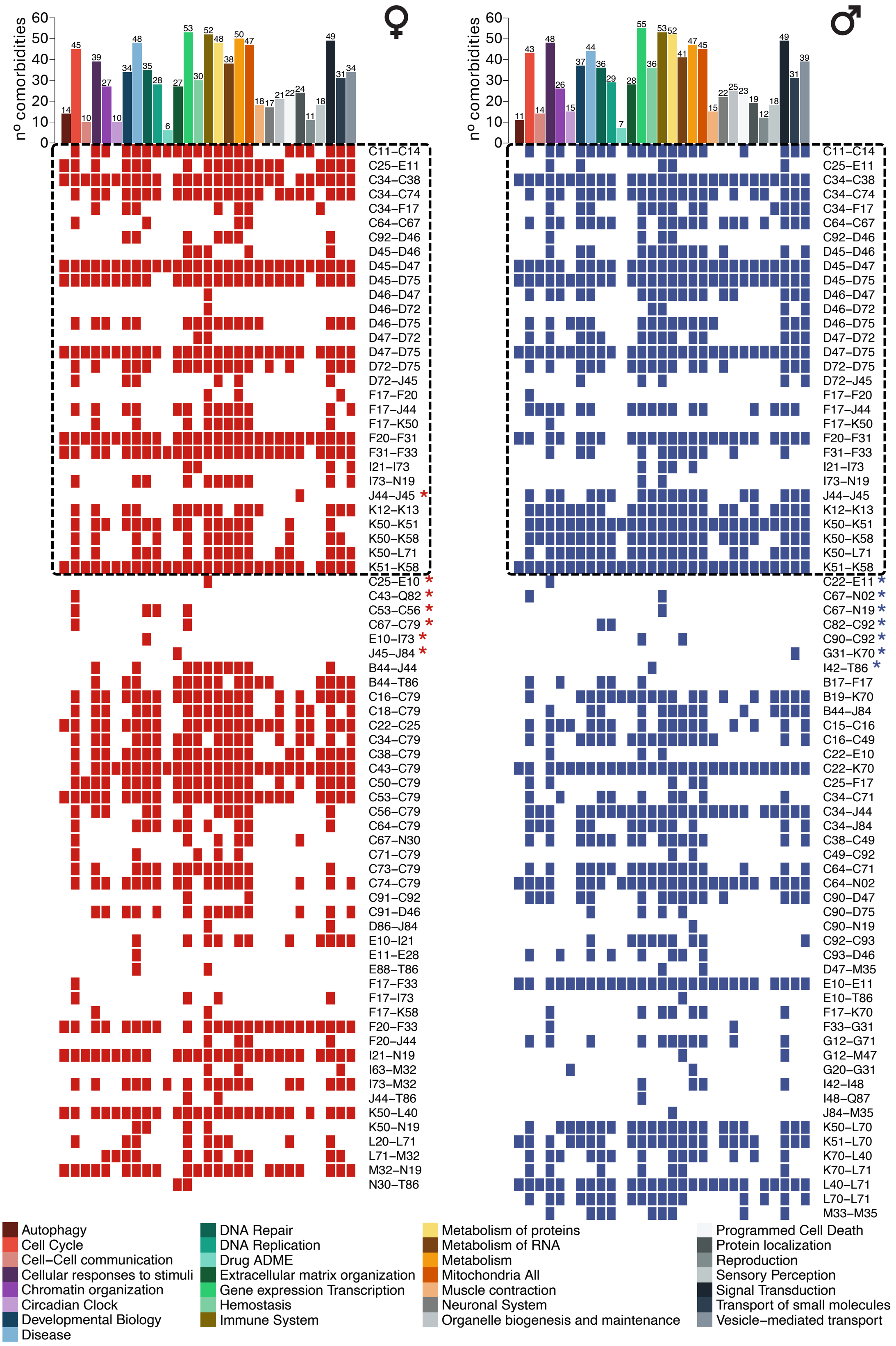


**Supplementary Figure 13. Comorbidities recovered by pathway category-specific networks.** Heatmap indicating in rows the comorbidities recovered in women (red) and men (blue) by generating disease networks based on their transcriptional similarity using only genes from the different Reactome categories and genes associated with mitochondrial processes (columns). Dashed Square highlights comorbidities identified in men and women. Asterisks denote comorbidities that are not detected when working with the entire set of genes.
